# A new genome-scale model enables prediction of cancer metabolic dependencies

**DOI:** 10.64898/2026.06.30.735578

**Authors:** Hoang V. Dinh, Asimina Zoitou, Jiazhen Zhang, Yihui Shen

## Abstract

Cancer cells rewire metabolism to support proliferation. Intriguingly, divergent metabolic choices are made to attain this common goal. Identifying the unique metabolic requirements for a specific cell has profound implications for cancer biology and precision medicine. Genome-scale metabolic models (GEMs) have emerged as powerful tools to systematically characterize, understand, and predict metabolism of cells and tissues. Despite being comprehensive, the current GEMs remain limited in their predictive power. Here, we present a new GEM of human cells, *in silico* <u>H</u>uman <u>M</u>etabolic <u>E</u>ssentiality (*i*HME), that significantly improves the prediction of metabolic dependencies at a reduced computational cost. Wse rationally downsized, curated, and corrected previous models to remove unsupported metabolic redundancies, which led to a slim model containing 4,377 reactions, 3,241 metabolites, and 1,825 genes. When used to reconstruct metabolic networks of 1,103 cancer cell lines, *i*HME recalled on average 84.6% of experimental essential genes, which is two-fold increase over previous models. Cholesterol biosynthesis was revealed to be the most reliably predicted pathway with alternative dependencies. Finally, we applied the model to reconstruct individualized networks and predict essential gene profiles for 8,384 patient tumor samples. Glucose transporter SLC2A1 (GLUT1) was identified as a context-specific dependency for head and neck cancers and ovarian cancer. Likewise, CDP-diacylglycerol synthase CDS2 was identified for skin cancer. Overall, *i*HME is a new genome-scale model for prediction of metabolic dependency at higher accuracy and computational efficiency.

## Introduction

Cancers display widespread changes in genes expression that support their deregulated growth. A common theme in these programs is metabolic rewiring that sustains proliferative needs. Altered metabolism often fulfills critical tasks such as taking up nutrients, obtaining energy, combating oxidative stress, and producing biomass, and therefore is recognized as a hallmark of cancer^1–3^.

More importantly, cancer metabolism can be exploited as a targetable vulnerability^4^. If a metabolic reaction is critical for any of the proliferative tasks, its disruption is very likely to impair cancer cell growth. Following this idea, inhibitors of nucleotide metabolism, such as antifolates, have achieved continued success in the clinics^5^. More recently, efforts to identify cancer-specific metabolic dependencies has led to an expanding panel of anti-cancer drugs^4,6–8^.

Cancers are molecularly diverse. Each cancer expresses a subset of metabolic genes, and thus is a unique context where metabolic vulnerability should be defined. Large-scale molecular profiling across thousands of cell lines and patient tumor tissues revealed substantial heterogeneity in global gene expression^9,10^. Importantly, cancer molecular classification is not exclusively driven by a single defining feature, but it is rather determined by combinatorial factors including tissue of origin, cell of origin, oncogene mutation, and microenvironment^10^. Therefore, despite that proliferation shares similar formula, the underlying metabolic networks appear very heterogeneous across cell lines and tumor tissues^11–13^. Indeed, the role of a metabolic gene in supporting proliferation diverges among different cells^12,14,15^. A particular challenge in defining cancer metabolic vulnerability is the multi-level redundancies encoded in human metabolic network: An enzyme’s function can be executed by isozymes or complexes encoded in different genes, a metabolic task can be fulfilled by pathways encompassing different sets of enzyme reactions, and a growth-essential metabolite can be obtained through intracellular conversion or from the environment. These fail-safe mechanisms may or may not exist in specific cellular contexts. Therefore, the effectiveness of blocking a metabolic reaction is uniquely determined by the active metabolic network in a cell.

Characterizing and targeting cancer metabolism could benefit from a systems approach that identifies context-specific metabolic dependency. Genome-wide screenings by RNA interference or CRISPR-Cas9 knock out provide powerful experimental approaches for systemically mapping the gene dependency landscape^16,17^. Several consortia efforts, including the most recent Project DepMap, have created a cancer dependency compendium across thousands of cancer cells^14,16–21^. Yet, experimental approaches are limited to contexts that can be modeled in labs. Computational approaches provide a scalable alternative in disease relevant contexts, such as patient tumors. Towards this, genome-scale metabolic models (GEM) enable integration of molecular data to simulate mechanistic outcomes of a metabolic network^22–25^. GEMs are mathematical representations of the entire metabolic network, explicitly accounting for genes, metabolites, and reaction stoichiometry from extensively curated biochemical knowledgebases^26–29^. Starting from a generic model, such as Human1^ref. 29^ and Recon3D^28^, context-specific models can be constructed using gene expression data of specific cells or tissues, and provide either structural analysis of metabolic networks^13,30,31^, or quantitative analysis of metabolic capacities across different cells or tissues using flux balance analysis (FBA)^32–34^. With the integration of patient-specific data, GEMS could identify novel disease biomarkers and provide insights for tailored therapeutic interventions^35–39^. However, despite the communities’ efforts to expand the generic human model^22,26–31,40^ and modeling algorithms and pipelines^41–45^, the predictive power of GEMs for gene perturbation remains modest – with only about 30-40% essential genes being correctly predicted^25,29,42^.

To facilitate the use of GEM for predicting metabolic dependency, we present a new GEM of human cells, *in silico* Human Metabolic Essentiality *(i*HME), that is rationally downsized, curated, and corrected from previous models. Using *i*HME as a base model, we derived cell-specific metabolic models from RNA-sequencing (RNA-seq) profiles of 1,103 human cancer cell lines. Together with a new set of proliferative metabolic tasks, our model predicted on average 84.6% of experimental essential genes, a 2.2 to 2.8-fold improvement from the previous models^25,29^. We also recovered 81% of lethal drugs that target metabolic genes in cell lines. We then applied *i*HME to generate metabolic models and predict essential genes for 8,384 human patient tumors. Intersecting the model predictions with genes upregulated in tumor compared to normal tissues, we identified SLC2A1 (GLUT1) as a context-specific dependency of head and neck cancer and ovarian cancers. Likewise, CDS2 was identified for skin cancers. Our findings in tumors were supported by screening in cell lines as well as literature-reported positive results from preclinical testing.

## Results

### Reconstruction of base genome-scale model

We first established a framework to reconstruct context-specific metabolic networks and predict gene essentiality. Our framework starts with a comprehensive base model, then applies the Integrative Network Inference for Tissues (INIT)^30^ algorithm to build context-specific models guided by transcript levels^46^, then further constrains the network with environmental nutrient availability, and finally evaluates proliferation-essential metabolic tasks using FBA^32^ (Fig. 1a). INIT eliminates reactions that are linked to lowly expressed genes if their neighboring reactions are not associated with highly expressed genes. By examining the functions of experimental essential genes (DepMap score < −0.5 in any cell line)^46^, we formulated a new set of 88 metabolic tasks to serve as biologically relevant optimization objectives, including biomass precursors, cofactors and prosthetic groups, energy and antioxidant maintenance, and protein modifications (see Methods, Supplementary Fig. S1a and S1b, and Supplementary Table S1). These tasks are the end goals of a metabolic network, drawn from cumulative knowledge of proliferative metabolism^27–29,36,47,48^. The new task panel preserves and significantly expands the previously used core tasks^30^ (Supplementary Fig. S1c).

**Figure 1.**
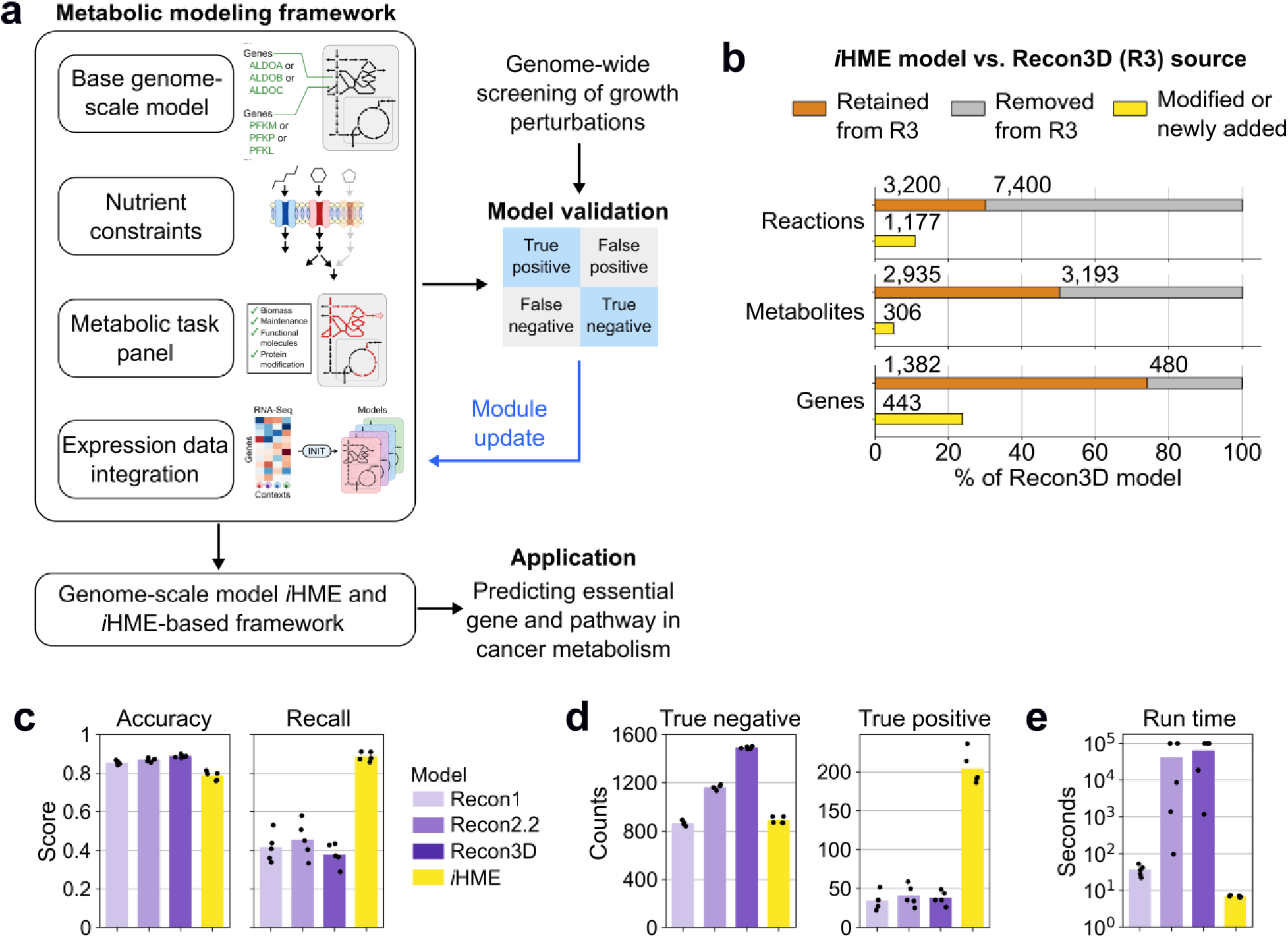
Genome-scale model *i*HME model and framework improve essential gene predictions. (**a**) Modeling framework setup and validation workflow. The four modules were systematically built to improve essential gene predictions, resulting in the refined *i*HME model and framework. (**b**) Comparison of *i*HME vs. Recon3D models. (**c,d,e**) Statistics across different base models (Recon and *i*HME models) and example cell lines (n = 5). (**c**) Accuracy and recall scores. (**d**) Counts of true negative (non-essential) and positive predictions (essential). (**e**) Run time (in seconds) for context-specific model reconstruction with INIT algorithm.

We then reconstructed metabolic models for five cell lines using Recon models as the base model^22,28,49^, and tested against experimental data of genome-wide loss-of-function screens from DepMap^14,46^, using a cutoff gene score of −0.5 for essentiality^21^ (Supplementary Fig. S2a). A gene was declared essential *in silico* if flux to any of the metabolic tasks is blocked following removal of the gene (Supplementary Fig. S2b). The results were then used to evaluate overall accuracy and recall of essential genes for each cell line. The models achieved a good accuracy (88.7%) and an improved recall of essential genes compared to an earlier report with the same models^29^ (Fig. 1c), indicating that the new task panel more comprehensively described the proliferative needs of cancer cells. Yet the recall score (37.5%) remained unsatisfactory throughout the three generations of Recon models. The high accuracy was rather driven by the dominant presence of non-essential genes^46^ (82.5% non-essential compared to 17.5% essential, Supplementary Fig. S2a) and true negative predictions (Fig. 1d). We reasoned that poor essentiality prediction was largely attributed to unverified bypass pathways – Recon3D contains 10,600 reactions, averaging to 5.69 reactions per gene (Fig. 1b), with 2,860 not associated with any genes. The redundancy in these models not only compromises recall but also greatly raises computational cost (Fig. 1e).

To improve the prediction of essential genes, we revised the base model to remove reaction redundancies. Starting with Recon3D model^28^ supplemented with updates from Human1 model^29^, we devised a data-guided iterative workflow to carefully downsize and re-curate the model (Fig. 1a and Supplementary Fig. S3, also see Methods). Briefly, we first curated biochemical reactions and their experimentally linked gene association available from databases including KEGG^50^, UniProt^51^, and HumanCyc^52^. Per iteration, we performed FBA analysis and removed unsupported metabolic redundancies if a reaction carries thermodynamically infeasible cycling flux^53^ or bypass flux for an essential gene^54^. Our final model, which we named *i*HME, contains 4,377 reactions, 3,241 metabolites, and 1,825 genes (Fig. 1b and Supplementary Table 2-4). Compared to Recon3D model, *i*HME has a much-reduced network size yet similar number of genes, averaging to 2.40 reactions per gene and containing fewer non-gene-associated reactions (906 reactions). When applied to construct context-specific models for five cancer cell lines, *i*HME achieved two-fold improvement in recall with six-fold more essential genes predicted (Fig. 1, c and d). Meanwhile, *i*HME effectively reduced the size of the active network (Supplementary Fig. S4), and greatly reduced the computational cost by 10^4^-fold (Fig. 1e).

### Genome-wide in silico validation of metabolic gene essentiality in cancer cell lines

To evaluate the predictive power of *i*HME, we extensively tested it in 1,103 cancer cell lines that have both genome-wide screens and gene expression data available in DepMap^46^ (Fig. 2a). A total of 1,103 context-specific models were first generated using *i*HME as the base model, forming an *in vitro* cancer metabolic reactome. While the pan-cancer model comprises 1,273 reactions that are active in at least one cell line, only about a half of these reactions are shared across >90% cell lines (Fig. 2b, core cancer, 661 reactions). For example, asparagine synthase is absent in 77 out of 91 lymphoid cancer cell lines, consistent with lymphoid cancer’s susceptibility to asparagine depletion^55^. On average, context-specific models for cancer cell lines contain 771 ± 30 reactions and 670 ± 19 genes.

**Figure 2.**
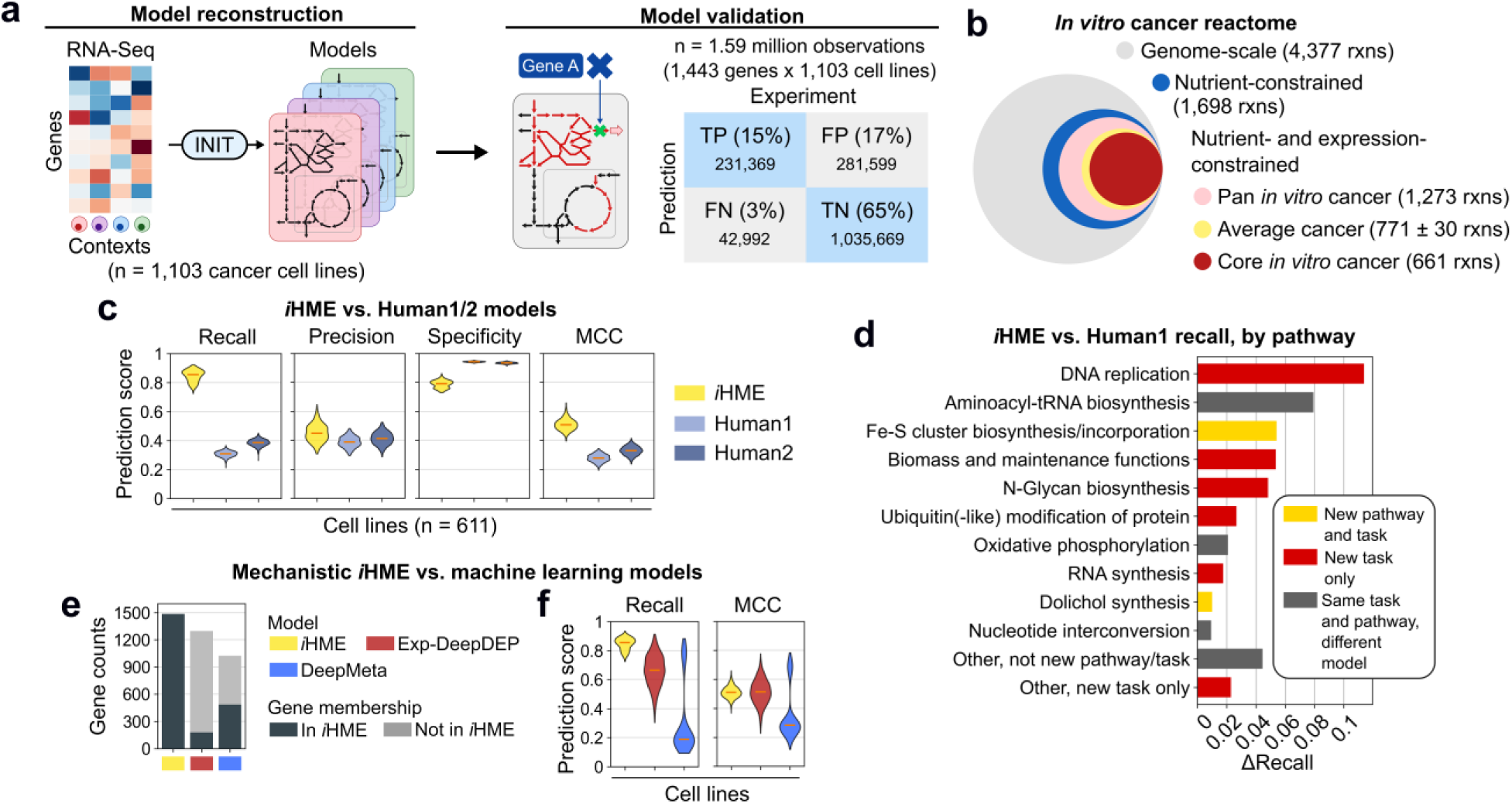
Reconstruction and validation of essential genes in cancer cell lines. (**a**) Reconstruction and validation workflow schematics. TP: True positive, FP: False positive, TN: True negative, FN: False negative. (**b**) The relative size of *in vitro* cancer reactome. “Core”: reactions present in 90% of cell lines. “Pan”: reactions in at least a cell line. (**c**) Comparison of prediction performance between *i*HME, Human1, and Human2 models. Violin plot displays distribution and median (orange line) of scores across overlapping cell lines (n = 611). (**d**) Pathway-specific recall difference between *i*HME and Human1 models. Annotations of new pathway and/or new task are color-coded. (**e,f**) Comparison of mechanistic *i*HME vs. machine learning models in terms of (**e**) gene coverage and (**f**) prediction scores of all cell lines. Violin plot displays distribution and median (orange line) of scores for cell lines, n = 1,103 for *i*HME, 1,103 for Exp-DeepDEP, and 784 for DeepMeta due to its data selection

Next, *in silico* knockout was performed for a total of 1,443 metabolic genes for each context-specific model, and cross-validated with experimental dependency (applying a cutoff gene score of −0.5 for essentiality^21^). The results were summarized in the 2-by-2 confusion matrix, which was then used to evaluate model performance (i.e., recall, precision, and specificity scores) for each cell line. Out of a total of 1.57 million predictions, 14.5% were true positive (TP), 65.1% true negative (TN), 17.7% false positive (FP), and 2.7% false negative (FN). This led to a mean recall score of 84.6% across 1,103 cell lines. Notably, compared to previous context-specific models built off Human1^ref. 29^, our models show a much-improved recall (2.8-fold), a comparable precision (1.2-fold), and slight drop in specificity (0.83-fold) for 611 overlapping cell lines (Fig. 2c). We then evaluated the overall performance with a balanced metric, Mathew’s Correlation Coefficient (MCC), that is computed from all prediction outcomes (TP, TN, FP, FN). An MCC score of 1 means perfect prediction while 0 is equivalent to random prediction. The new models scored significantly better (MCC = 0.51) than the previous models (MCC = 0.28) (Fig. 2c). The better recall was largely contributed by newly included metabolic tasks, and to a lesser extent by new pathways and revised gene-reaction rules (Fig. 2d). We additionally validated model predictions of pharmacological inhibitors of metabolism from the PRISM secondary drug screening dataset^6^ (Supplementary Fig. S5a-b). *i*HME recalled 81.3% of lethal drug administration cases, 1.8-fold more than what was simply inferred from gene dependency, likely due to the capability of metabolic models to capture reaction dependency in addition to gene dependency (Supplementary Fig. S5c-f).

We noted that precision (TP/(TP+FP)) remains moderate regardless of base models used. Interestingly, model predictions of TP and FP are pathway dependent (Supplementary Fig. S6). Among the 72 metabolic pathways, TP predictions are enriched in macromolecule synthesis, ROS detoxification, and Fe-S synthesis. In contrast, FP predictions are enriched in enzyme cofactor synthesis (Supplementary Fig. S6). As cofactors are essential for enzyme function, these false positives suggest the existence of adaptive mechanisms that remain unpredictable. These mechanisms have been actively explored in case studies for NADP^+ 56,57^, glutathione^58^, heme^59^, and flavin^60^ cofactors, to explain differential sensitivity in cells.

To explore if cellular context contains hidden correlations that elude our metabolic network, we further benchmarked our knowledge-driven *i*HME with emerging data-driven machine learning (ML) models (Fig. 2e-f). Using the same gene expression input data, we compared our model’s performance to those from two deep learning models, Exp-DeepDEP^61^ and DeepMeta^62^. Overall, ML models make predictions for a smaller set of metabolic genes (Fig. 2e), as only genes with highly varied gene scores were considered^61,62^. Moreover, ML models include non-metabolic genes in their input, which link external genes to metabolism. Remarkably, *i*HME–derived mechanistic models on average have the highest recall and comparable or higher MCC scores for metabolic genes comparing to either ML model (Fig. 2f). *i*HME displayed similar recall even for the smaller subset of genes predicted by ML models (Supplementary Fig. S7). Therefore, our mechanistic modeling approach provides a greater coverage of metabolic genes and performed similarly to if not better than ML models, suggesting that metabolic dependencies are mostly contained in the metabolic network.

In summary, we reconstructed a new human metabolic model, *i*HME, that is slim yet extensively curated. Combined with a newly crafted metabolic task panel, *i*HME allows rapid construction of context specific models that better identify essential metabolic genes.

### Context-specific metabolic dependencies of cancer cell lines

Inherent to its network structure, an advantage of metabolic models is their ability to provide pathway insights. As the human metabolic network has built-in redundancies at the alternative isoform (isozyme) and pathway levels, we then asked what metabolic genes are uniquely needed in subsets of cell types, and what explains their variable dependencies. To answer these, we first evaluated prediction from the base *i*HME, nutrient-constrained base model, or the pan-cancer model containing the reactions from all the context-specific models. Compared to these generic context-free models, context-specific models show significantly better recall (Fig. 3a), indicating that gene expression largely shaped the metabolic dependencies of different cancer cell lines.

**Figure 3.**
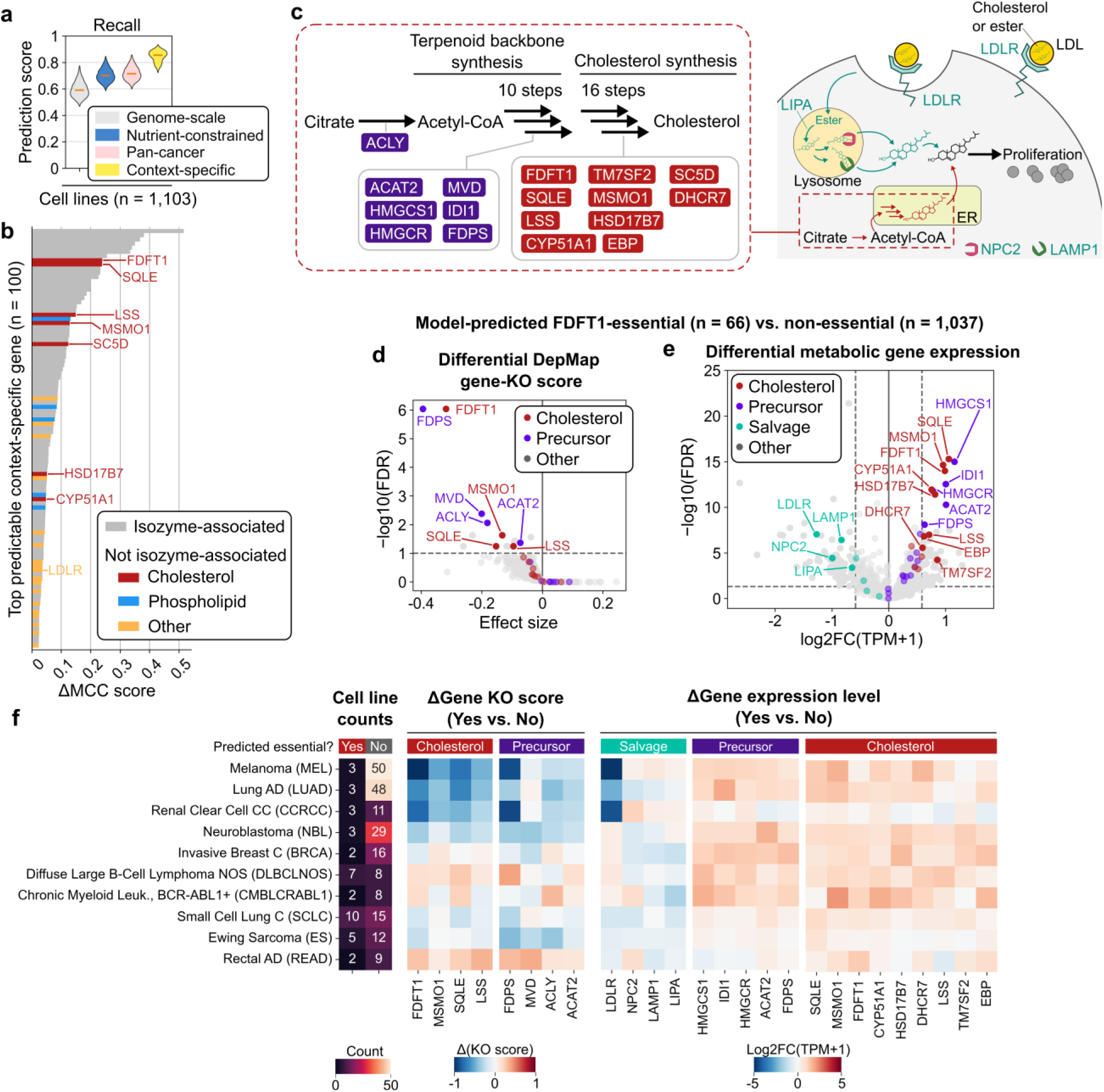
Context-specific models predict cellular dependencies on isozymes and alternative pathways. (**a**) Essential gene recall scores from context-specific and context-free models. Violin plot displays distribution and median (orange line) across cell lines (n = 1,103). (**b**) MCC score difference between context-specific and nutrient-constrained models, for top 100 genes. (**c**) Cholesterol synthesis and salvage pathways visualization. (**d,e,f**) Comparison of DepMap gene score and gene expression between model-predicted FDFT1-essential (n = 66) vs. non-essential (n = 1,037) cell lines, for n = 1,430 metabolic genes. (**d**) Differential DepMap gene knockout score analysis, where negative effect size indicates more severe growth reduction with gene knockout. The cut-off (dashed lines) is FDR = 0.1 from one-tail t-test. (**e**) Differential gene expression analysis. Cut-offs (dashed lines) are: fold-change = 1.5 and FDR = 0.1 from two-tail t-test. (**f**) Gene knockout score and gene expression level differences for cell lines grouped according to Oncotree disease classifications. Cancer diseases with n = 1 for predicted essential are not shown.

We then reasoned that expression-driven gene dependencies could be identified by an increase of a gene’s MCC score between models with or without the gene expression context (Fig. 3b, Supplementary Table 5). Within the top 100 genes with MCC score increase, many are associated with isozymes (Fig. 3b). These include subunits of V-type ATPase (ATP6V) for intracellular acidification, PIP5K1A for phosphatidylinositol 4,5-bisphosphate synthesis, and IMPDH2 for purine synthesis. Model predictions of essentiality are driven not only by the expression but also from the lack of its alternative genes (Supplementary Fig. S8a). Specifically, while all the three PIP5K isozymes are present in normal tissues, the expression is skewed towards PIP5K1A in cancer cell lines, leading to its predicted essentiality in about two thirds of the cell lines (Supplementary Fig. S8b-d). Similarly, IMPDH2 essentiality is mostly driven by the absence of IMPDH1 in about half of the cell lines (Supplementary Fig. S8e-g). Our analysis thus identifies the cellular contexts that may respond to inhibition of the above isozymes^63–65^.

Interestingly, we also found a few context dependencies without isozyme association. These include genes in cholesterol biosynthesis and phospholipid metabolism (Fig. 3b). Cholesterol is an essential component of cell membranes, and regulates membrane fluidity and permeability^66^. Cholesterol is synthesized through 10 reactions that convert acetyl-CoA to the terpenoid backbone followed by 16 reactions committed to cholesterol (Fig. 3c). Among these, FDFT1 and SQLE, which catalyze the squalene synthesis and epoxidation, are ranked as top genes predicted to be context-specific essentials. Out of the 1,103 cell lines, a total of 66 are predicted to depend on cholesterol synthesis, which indeed showed more negative growth impact from the loss of cholesterol synthesis or precursor supply in experiments (Fig. 3d). We then searched for alternative pathways in the cell lines predicted to be non-dependent, and found that cholesterol flux is instead fulfilled by the salvage pathway via lipoprotein uptake (LDLR) and lysosome processing (NPC2, LAMP1, LIPA)^67^ (Fig. 3c). Differential gene expression analysis confirmed that the model prediction is driven by a concerted upregulation of genes for *de novo* synthesis and down regulation of genes for salvage (Fig. 3e).

Importantly, the requirement of cholesterol biosynthesis is not associated with tissue of origin or disease subtypes obtained from Oncotree classifications^68^. Rather, it is strongly predicted in subsets of cell types across Melanoma (MEL), Lung Adenocarcinoma (LUAD), Renal Clear Cell Carcinoma (CCRCC), and Neuroblastoma (NBL) (Fig. 3f). This suggests that cholesterol synthesis is a common metabolic adaptation in cancers. Consistently, we found that loss of SREBP2, a transcriptional factor that stimulates cholesterol synthesis in response to cholesterol deficiency^69^, also had a small but significant negative impact in cells dependent on cholesterol synthesis (Supplementary Fig. S9). In summary, the cholesterol procurement through *de novo* synthesis or salvage is a predictable metabolic vulnerability independent of cancer classification. This suggests that inhibiting cholesterol synthesis, such as with statins^70,71^, can be tailored for case-specific cancers.

### Reconstruction of metabolic networks for human tumors

To systemically explore the metabolic heterogeneity among human tumors, we applied our pipeline to reconstruct tailored metabolic networks for a total of 8,384 patient tumors across 17 organs that have been characterized by The Cancer Genome Atlas (TCGA)^10^ and Human Pathology Atlas (HPA)^13,72^ projects (Fig. 4a and 4b). The RNA-seq data, available in the HPA data repository^13,72^, were used to construct context-specific models, which were then constrained by human plasma nutrient availability to recapitulate the *in vivo* context^73,74^. From these models we performed 12.4 million *in silico* gene deletions, 33% of which are predicted essential (Fig. 4a). At an overview, the tumor reactome consists of 1,378 pan reactions (i.e., present in at least a tumor sample) and 594 core reactions (i.e., present in >90% of tumor samples), averaging to 760 ± 63 reactions in each network (Fig. 4c). Therefore, the *in vivo* tumor metabolic networks appear more variable compared to *in vitro* cell lines (Fig. 2b). Among organ systems, esophagus/stomach tumors present the smallest conserved network of 597 reactions (present in >90% of the samples), whereas liver tumors have the largest network consisting of 651 conserved reactions. To examine the metabolic characteristics of human tumors, we then clustered the tumors based on the three model-derived features: (i) gene presence, (ii) reaction presence, and (iii) gene essentiality (Fig. 4b). The clusters largely reflect organ systems (Fig. 4d), consistent with tissue of origin driving the molecular diversity of tumors^10^.

**Figure 4.**
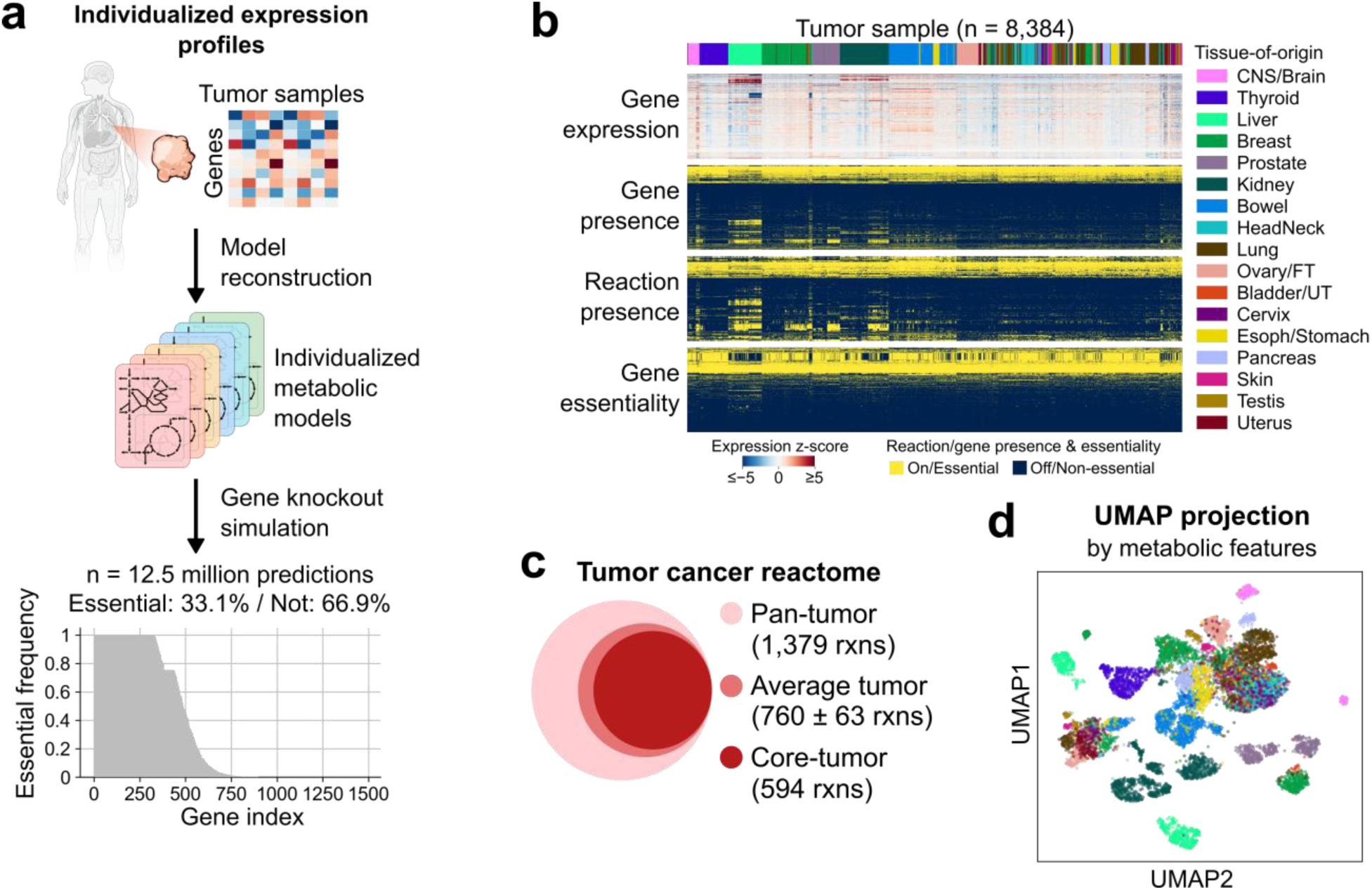
Metabolic reconstruction and metabolic feature analysis of human tumors. (**a**) Workflow of model reconstruction and gene knockout analysis for human tumors. Bottom panel shows distribution of essentiality frequency across tumor samples for all 1,479 genes. (**b**) Overview of gene expression data input and metabolic feature outputs (gene presence, reaction presence, and gene essentiality) for tumor samples (n = 8,384). Tumors’ tissue-of-origins are color-coded. Row features that are uniformly zero or one are not shown. (**c**) Relative size of tumor reactome. Pan-reactions are present in at least one sample. Core-reactions are present in > 90% of samples. (**d**) Uniform manifold approximation and projection (UMAP) plot of tumor samples using model-derived metabolic features. Tumors’ tissue-of-origins are color-coded as in (b).

### Priority genes for tumor metabolism by tissue of origin

We next sought to define priority genes of tumor metabolism for each TCGA tumor subtype. A gene is classified as a priority gene based on the two criteria: (i) two-fold upregulated expression compared to normal tissue (FDR < 0.05, see Methods for details on expression data collection and statistical test), and (ii) being predicted essential in more than 75% samples (i.e., essentiality frequency > 0.75) (Fig. 5a). Out of 29,113 pairs of genes and tumor types, 2,237 pairs are upregulated, among which 561 pairs are also frequently essential (Supplementary Table 6). A priority list of 226 genes was obtained, representing pathways including oxidative phosphorylation, RNA synthesis, DNA replication, N-glycan synthesis, amino acyl-tRNA charging, Fe-S cluster biosynthesis, and nucleotide interconversion (Fig. 5b). Looking at essential frequency distribution across 20 TCGA tumor subtypes, 167 out of 226 priority genes are predicted commonly essential (in at least 18 out of 20 subtypes), which indeed significantly overlap with common essential genes in DepMap screenings^46^ (104 out of 167, 62.3%).

**Figure 5.**
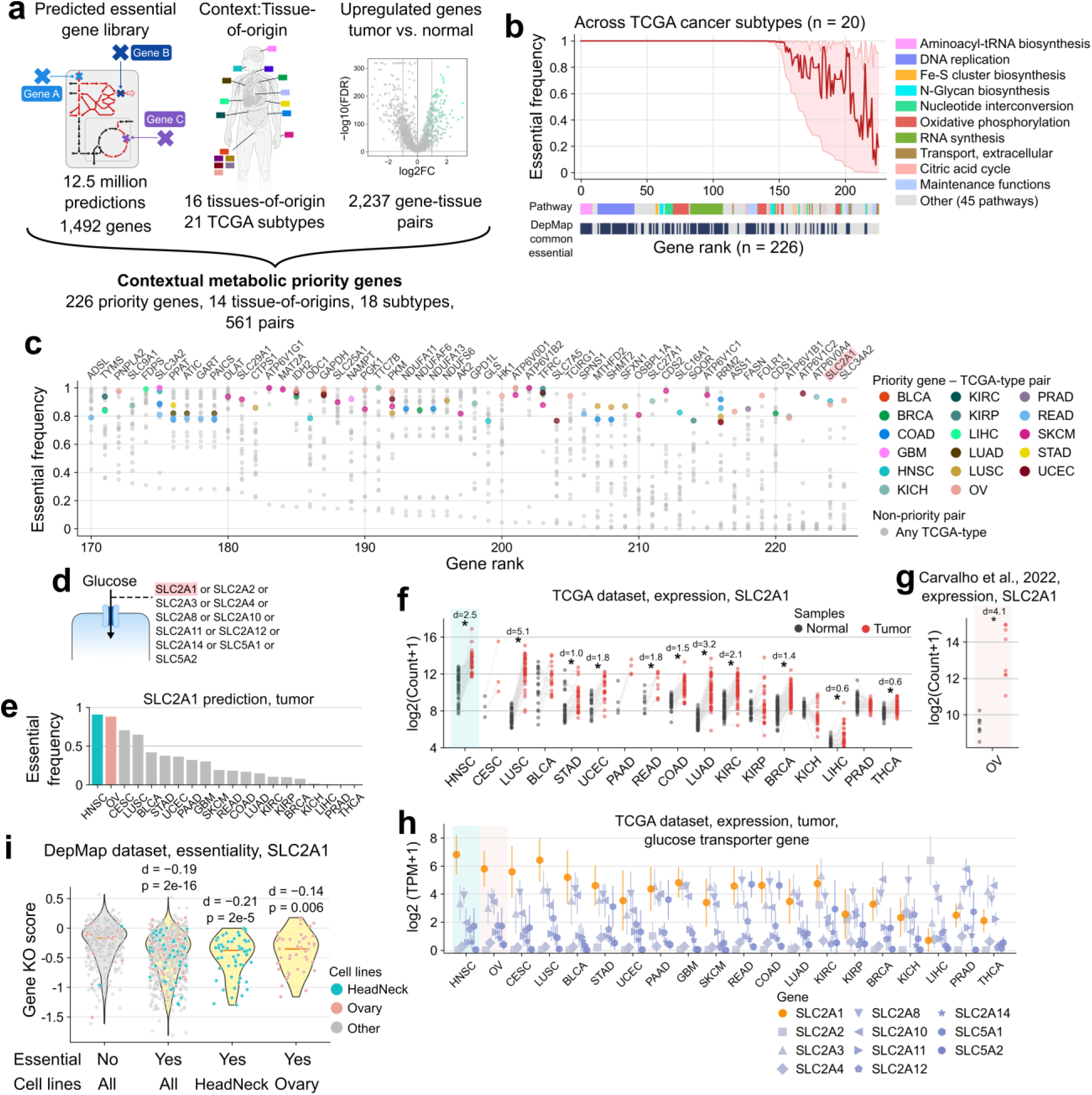
Identifying priority genes in tumor metabolism. (**a**) Criteria for identifying priority genes. They are specific to tumor subtypes defined by TCGA, significantly upregulated, and essential in model simulation. (**b**) Essential frequency across TCGA subtypes for priority genes (n = 226). Metabolic pathway and DepMap common essential annotations for genes are provided. Genes are ranked by minimum essential frequency in descending order. Upper, middle, and lower marks are maximum, average, and minimum essential frequency across TCGA subtypes. (**c**) Essential frequency for the bottom quarter of ranked list of genes in (**b**). Scatter points track the essential frequency value of gene – TCGA-subtype pairs. Genes are tracked on x-axis and TCGA-subtypes are tracked by colors. (**d**) Genes annotated to be glucose transporters. (**e**) Essential frequency of SLC2A1 in different TCGA tumor subtypes. (**f**) Expression level of SLC2A1 in paired tumor and tumor-adjacent tissue samples from TCGA database. Expression is in log2(Count+1). Log2-fold change (d-value) and significance (*, if FDR < 0.05) from paired samples t-test are provided. n = 3 to 98 paired samples across TCGA tumor subtypes. Head-Neck Squamous Cell Carcinoma (HNSC) is highlighted. (**g**) Expression level of SLC2A1 in ovarian (OV) tumor and normal tissue samples from Carvalho et al., 2022^ref. 78^. Description is the same with (**f**) except for the usage of unpaired two-tail t-test. n = 8 and 8 for unpaired tumor and normal tissue samples, respectively. (**h**) Expression level of glucose transporters in tumor. Expression is in log2(TPM+1). Mean and standard deviation values (across samples) are shown. n = 64 to 1,022 samples across TCGA tumor subtypes. HNSC and OV are highlighted. (**i**) DepMap gene knockout scores of SLC2A1 across cell lines. Distributions of scores were plotted for groups of cell lines predicted by the model to be essential or non-essential and annotated to be from specific tissue of origins. One-tail t-tests were performed to compare scores of essential to those of non-essential groups.

Common essential genes are universal bottlenecks in cellular proliferation, yet conditionally essential genes allow more tumor specificity when therapeutically targeted. Among the priority gene list, there are 55 genes showing variable essentiality frequency across tissue contexts (i.e., lowest tissue essential frequency being below 0.4) (Fig. 5c). Remarkably, among them, we recovered many metabolic targets that have proceeded to clinical trials, including FASN, RRM2, SLC16A1, SHMT2, SLC7A5, GLS, complex I, NAMPT, ODC1, IDH2, and TYMS^8^. Our analysis highlights the need to consider and validate the metabolic context in targeted therapy. For example, SLC2A1, which encodes a facilitated glucose transporter GLUT1 (Fig. 5d), is predicted to be uniquely required in head and neck tumors and ovarian tumors (Fig. 5e). This prediction is supported by significant upregulation of SLC2A1 in tumor tissues compared to their normal counterparts (Fig. 5f and g), as well as favorable expression of SLC2A1 over other glucose transporters (Fig. 5h). The tumor specificity is further supported by significant dependence on SLC2A1 in head and neck cell lines and ovarian cell lines (p = 2e-5 for head and neck, and p = 0.006 for ovarian) (Fig. 5i).

The conditionally essential list also recovered isozymes of CDP-diacylglycerol synthase, CDS1 and CDS2, which are needed for the synthesis of the essential signaling molecule phosphatidylinositol (3,4,5)-triphosphate (PIP_3_) (Supplementary Fig. S10). CDS2 is uniquely important to skin tumors due to its upregulation compared to normal skin, its high essential frequency, and its favorable expression over CDS1 in skin tumors (Supplementary Fig. S10). Indeed, experimental data in cancer cell lines show that skin cancer cell lines show significant lower gene score for CDS2 than non-dependent cell lines (p = 6e-18).

Importantly, pharmacological inhibition of SLC2A1 has shown efficacy in head and neck cancer cell lines and mouse models^75,76^. CDS1 and CDS2 have been validated as synthetically lethal pair in mesenchymal-like cancers^77^. Together, they support that our systematic analysis reveals not only meaningful priority genes, but also the tissue metabolic context that supports their essential role.

## Discussion

We curated a new genome-scale metabolic (GEM) model of human metabolism, *i*HME model, to predict metabolic dependency. On the basis of previous models^28,29,40^, thousands of manual curations were made to reactions and genes in the *i*HME model to improve the recall of essential genes. We rationally reduced the size of the model, removing unverified redundancy while greatly enhancing the computational efficiency. This permitted a rapid iteration of model curation to reconcile model prediction and experimental data, achieving a higher recall rate than the most updated Human2 model^25^ (i.e., 84.6% vs. 38.5%). We also noted that our software implementation of INIT algorithm has an extension, which is a post-optimization module to adjust gene-reaction rules, that could improve the recall rate for isozymes. In addition, *i*HME model is equipped with an extended metabolic task panel that more comprehensively model the needs of proliferation. Altogether, these technical improvements made substantial improvement to prediction power and will also provide a computationally efficient platform for genome-scale metabolic modeling.

Constructing context-specific metabolic models from *i*HME reveals alternative modalities used by human cells to achieve certain metabolic tasks. For example, on the pathway level, cholesterol demand is fulfilled by cholesterol synthesis and salvage; on the reaction level, alternative transporters or isozymes, such as glucose transporters and CDP-diacylglycerol synthase isozymes, confer redundancy or synthetic lethality. A subset of predicted metabolic dependencies has been associated to contexts, for example, SLC2A1 to head and neck and ovary tumors, and CDS2 to skin tumor. In human tumors, we proposed 226 metabolic dependencies and their associated contexts that could be worthy of further experimental validation.

Despite exhaustive efforts in reconstructing network and gene-reaction rules, false positives in essential gene prediction remain mostly in cofactor synthesis pathways. It is possible that the experimental time course used in the DepMap screening was not sufficient to deplete these cofactors. Alternatively, the assumption of cofactor demands being common essential metabolic tasks may be questioned. Unlike biomass components that are absolutely necessary for proliferation, cofactors are needed for enzyme functions. Several studies have discovered conditional essentiality of cofactor synthesis. For example, mitochondrial NADP^+^ synthesis is needed when proline is depleted^56^. Cytosolic NADP^+^ synthesis is needed under low-folate conditions to support nucleotide synthesis^57^. Glutathione synthesis is needed when loss of deubiquitinating functions aggravates proteotoxic and ER stress^58^. Heme synthesis is needed by Kras-driven cancer cells *in vivo* but not *in vitro*^59^. Flavin synthesis is needed to maintain nucleotide balance specifically in leukemia^60^. Overall, the requirement of cofactor synthesis appears to be conditional on cellular and environmental factors that are not completely understood. A novel modeling paradigm extending beyond flux balance will be required to computationally solve metabolic dependency that involves complex interactions, such as those between cofactors, metabolism, nutrients, non-metabolic stresses, and contexts.

A limitation in our approach was the use of only mRNA expression data to emulate active metabolic reactions. Reaction flux does not always correlate with mRNA level, and may also be regulated by translation, post-translational modification, and substrate or allosteric interaction. In addition, network approach cannot quantitatively assess the degree of impact from gene loss, such as the gene score in genome-wide screen. Under the assumption that genes carrying greater fluxes may pose a larger impact on metabolic fitness upon deletion, a semi-quantitative model incorporating metabolic measurement may be built in the future. Another limitation is the use of bulk expression data in our prediction of tumor metabolic dependency. The tumor microenvironment (TME) comprises various types of cells (e.g., cancer, immune, stromal, and vascular cells)^79^. In addition to cell types, tumor interstitial fluid has been measured to be substantially different in metabolite abundance compared to plasma^80^. Metabolite availability for cells and metabolite feeding between cells within TME can influence tumor malignancy and response to treatment^81–83^. This calls for a more faithful reconstruction of tumor microenvironment to account for cellular community, identify active fluxes in cell types, and recover meaningful cell-cell interactions from a large number of combinatorial interactions. For example, by performing *in vivo* ^13^C-metabolomics and computational metabolic flux analysis, Meghdadi and coworkers systematically resolved cell-cell interaction in serine metabolism and purine synthesis and salvage preference across cell types^82^. With the advance in single cell and spatial transcriptomics^84^ and in computational efficiency offered by *i*HME model, it is possible to analyze metabolic network, flux, and interactions across thousands of single cells recovered from the current technology. This development will be instrumental to solve any exceptional metabolic dependency that can be unmasked with considering single cells in TME.

## Methods

### *i*HME model reconstruction and curation

The starting Recon3D model was downloaded from the BiGG database (http://bigg.ucsd.edu/). Test simulations revealed many issues affecting model predictions. These issues were hereby mentioned and fixed in the order of their discoveries. First, nutrient sources introduced by reverse fluxes of 95 sink reactions, which are not exchange reactions, were eliminated (i.e., lower bounds set to zero). We then identified 2,655 reactions (i.e., with unbounded flux values) participating in thermodynamically infeasible flux cycles via flux variability analysis (FVA)^85^ with all exchange reactions closed. Also presented in the model are 3,160 metabolic and 1,502 transport reactions that are not associated with any genes (i.e., hypothetical reactions). Because fixing these unverified and/or erroneous reactions requires manual curation and the Recon3D model size is large, we first removed peripheral pathways before manual curation. Being removed are peripheral pathways metabolizing the following molecules: (i) drugs, (ii) peptides (excluding glutathione), (iii) bile acids, (iv) large complex carbohydrate-derived and protein-derived molecules (e.g., carbohydrates, proteins, glycoproteins, and glycolipids), (v) toxins, (vi) D-amino acids, and (vii) hormones and signaling molecules (excluding serotonin). Furthermore, to simplify lipid metabolism, lipid species with acyl groups of different chain lengths and degrees of saturation were replaced with species with a generic acyl group that can be specified. We estimated the acyl pool in the model to compose of observed major fatty acids (i.e., >1% of total fraction) in phosphatidylcholines of human blood samples^86^. The fatty acids are palmitic (C16:0) (at molar fraction of 34.17%), stearic (C18:0) (15.61%), oleic (C18:1 n-9) (11.03%), linoleic (C18:2 n-6) (23.91%), dihomo-gamma-linoleic (C20:3 n-6) (2.62%), arachidonic (C20:4 n-6) (9.06%), and cervonic (C22:6 n-3) (3.60%). Acyl groups in all lipid species are represented by a single average acyl group formed by the following reaction:

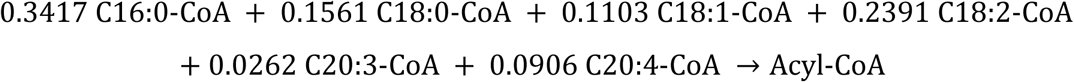

For central and central-adjacent metabolic pathways, we eliminated thermodynamically infeasible flux cycles by removing reactions that are hypothetical (i.e., not associated with enzymes) or do not match functional genomic and experimental evidence. For example, invalidated reactions associate with an unverified alternate cofactor (e.g., GTP instead of ATP) or perform hypothetical functions that were not substantiated. We consulted human-curated UniProt^51^ and HumanCyc^52^ databases as trusted collections of functional annotation and experimental evidence.

Transporters for the same metabolite can form flux cycles (i.e., constituted of separate uptake and secretion reactions) or create unresolved alternate flux distributions (i.e., between separate transport mechanisms such as facilitated diffusion and ion-coupled transportation). Thus, for each metabolite we replaced those transporter reactions with a single diffusion transport reaction since chemical gradient consideration is outside the purview of stoichiometric-only modeling. This replacement resolves transporter flux cycle and alternative flux issues.

Next, flux leakages associated with hypothetical and unverified reactions were identified and eliminated because they rescue loss of canonical pathways. To identify these hypothetical reactions, flux balance analysis (FBA)^32^ was performed under canonical pathway knocked out. Hypothetical reactions that carried flux and rescued *in silico* cells were removed one-by-one until flux simulation returned solutions with flux blocked or rescued by another canonical pathway.

Biomass reaction was adapted from Zielinski et al., 2017^ref. 87^ and then updated. Biomass DNA, RNA, and protein precursors were corrected (from dNMP, NMP, and amino acids) to dNTP, NTP, and amino acyl-tRNA, respectively. tRNA-synthetase reactions were added to the model. Biomass carbohydrate precursors were updated (from glycogen) to the monomer units of N-acetyl-D-glucosamine (GlcNAc), mannose (Man), glucose (Glc), galactose (Gal), fucose (Fuc), and N-acetylneuraminate (Neu5Ac), and N-acetyl-D-galactosamine (GalNAc). The molar composition of carbohydrate units was set to 11.75% GlcNAc, 28.01% Man, 13.78% Glc, 20.21% Gal, 1.24% Fuc, 15.50% Neu5Ac, and 10.25% GalNAc, which is a theoretical estimation based on glycoproteome and glycosphingolipid measurements and knowledge of glycosylation sites for CHO cells^88^. Compositions of biomass free fatty acids and acyl groups of biomass lipids were updated to the measurements^86^. ATP synthase mechanistic ATP/proton ratio was corrected to 3/8 based on enzyme structure observation^89^.

Being newly added to the model are the pathways of (i) iron storage in Ferritin, (ii) Fe-S synthesis and incorporation, and (iii) (for the most part) methyl- and adenosyl-cob(III)alamin (i.e., vitamin B12) synthesis. Fatty acid uptake and mitochondrial and peroxisomal fatty acid oxidation were reconstructed anew for the major fatty acids of palmitic (C16:0), stearic (C18:0), oleic (C18:1 n-9), linoleic (C18:2 n-6), dihomo-gamma-linoleic (C20:3 n-6), arachidonic (C20:4 n-6), and cervonic (C22:6 n-3) acids.

Finally, many changes for reactions and their gene-reaction rules were made so that the flux predictions of the model satisfy these criteria: (i) simulation using biomass reaction was supported by canonical central pathways (i.e., glycolysis, TCA cycle, and oxidative phosphorylation) and biosynthetic pathways (i.e., for amino acids, nucleotides, carbohydrates, and lipids) instead of by unverified alternative pathways (i.e., by unrelated pathways and by reactions without gene-associations); and (ii) simulation of gene knockout are consistent with DepMap gene knockout data.

Once the model had had its unverified redundant pathways trimmed off, the model went through iterations of simulation and validation with DepMap cell line data. False negatives and positives were exhaustively checked and resolved. False negatives were resolved by adding new metabolic tasks. False positives were resolved by identification of absent isozymes, absent alternative pathways, or false gene annotation. For example, cholesterol salvage pathway was searched for and added based on prior false positives in cholesterol synthesis pathway. False gene annotation comes from sequence homology-based annotation, such as the recently discovered false annotation of pantothenate kinase activity by PANK4 protein^90^. Discovery and removal of false associations to function (e.g., PANK4) in the model solved corresponding false positives.

The spreadsheet version of the model is in Supplementary Tables S2-4. Models in simulation-compatible formats (i.e., COBRApy-JSON and MATLAB) are in https://github.com/hvdinh16/iHME_model.

### Nutrient and secretory metabolites panel formulation

Flux balance analysis in this work is constrained to maximal flux capacities (except INIT algorithm stage 1, which is to be explained later). Nutrient uptake capacity cannot be unconstrained because in simulation any unconstrained nutrients (e.g., any amino acids) can replace the needs for major nutrients (e.g., glucose and glutamine), rendering model failed to simulate major nutrient catabolism. For a nutrient exchange reaction *j_EX_* ∈ {*Nutrient exchange set*}, the constraint is 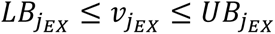, where maximal capacity is set in the lower bound 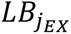 <u>in negative value</u> under exchange flux convention.

For nutrient recorded in the CORE profile^91^, the maximal nutrient uptake rate 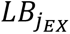 values were set to standard values derived from the maximally measured values for NCI-60 cancer cell lines^87,91^. The standard value 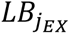 (for exchange reaction *j_EX_*) was estimated by the following equations:

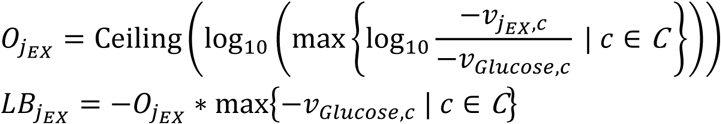

where 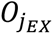 is the order of magnitude of maximal ratio of nutrient 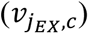 to glucose uptake (*v_Glucose_*_,*c*_) fluxes, and 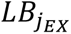 is the product of 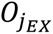 with the maximal glucose uptake rate (which is **−**5 mmol gDW^−1^ h^−1^). The experimental values were estimated across cell line *c* ∈ *C* (i.e., NCI-60 cell lines). Maximal uptake capacity 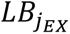 values were used in simulation. For an exchange reaction that is not in the nutrient panel, *j_EX_* ∈ {*All exchange set*} \ {*Nutrient exchange set*}, the constraint is 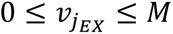.

For nutrient not recorded in the CORE profile ^91^, the maximal nutrient uptake rate 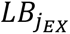 values were anchored to maximal glucose uptake rate 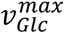 (i.e., −5 mmol gDW^−1^ h^−1^), maximal cancer cell line growth rate (i.e., 0.04 h^−1^), and biomass reaction coefficients, which are hereby listed:

(i) The sum of molar uptake flux of six fatty acids is set to 1/3 of 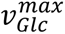, assuming that fatty acids can be major nutrients like glucose. The ratio 1/3 is derived from one divided by 18 carbons in an average fatty acid over 6 carbons in glucose. Individual fatty acid uptake rate is all set to 1/18 of glucose uptake, which is 1/6 (i.e., one out of six fatty acids) of 1/3 (i.e., a third of glucose uptake rate because the number of carbon of fatty acid is roughly three times that of glucose).
(ii) Histidine and cysteine uptake are individually set to 0.1 of 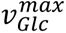, per other proteinogenic amino acid data.
(iii) Riboflavin (B2), biotin (B7), pyridoxin (B6), pantothenate (B5), thamine (B1), folate (B9), aquacob(II)alamin (B12), Fe^2+^, and Fe^3+^ uptake are individually set to 0.01 of 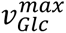, per nicotinamide (B3) data.
(iv) Nicotinamide D-ribonucleotide (NMN) (B3) and nicotinic acid (B3) uptake are individually set to 0.001, an order of magnitude lower than the value for nicotinamide (B3) data.
(v) Cholesterol and cholesterol ester uptake are individually set to 0.001524 mmol gDW^−1^ h^−1^ uptake flux, which is the product of maximal growth rate and the sum of biomass reaction coefficients for cholesterol and cholesterol ester.
(vi) Phosphatidylcholine (PC), lysophosphatidylcholine (LPC), phosphatidylethanolamine (PE), and lysophosphatidylethanolamine (LPE) uptake are individually set to 0.003492 mmol gDW^−1^ h^−1^, which is the product of maximal growth rate and the sum of biomass reaction coefficients for PC, LPC, PE, phosphatidylserine, phosphatidic acid, and cardiolipin (x2, only on cardiolipin).
(vii) Triacylglycerol uptake is set to 0.000127 mmol gDW^−1^ h^−1^, which is the product of maximal growth rate and the sum of biomass reaction coefficients for monoacylglycerol and diacylglycerol.
(viii) Lactate uptake is set to 0.1 of glucose uptake based on concentration ratio in plasma.
(ix) 3-Hydroxybutyrate, acetate, acetone, glycine betaine, formate, fructose, reduced glutathione, and pyruvate uptake are individually set to 0.01 of 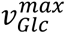 based on concentration ratio in plasma.
(x) The uptake rates of ambient molecules are set to arbitrary large number (i.e., **−**1e6 in this work). These are oxygen, phosphate, sulphate, proton, water, and CO_2_.

Secretion fluxes are turned off for all metabolites except for those that were observed in the CORE profile *in vitro*^91^ and in the serum metabolomics *in vivo*^73,74^. The secretion metabolite panels for in vitro and in vivo include 77 and 86 metabolites, respectively. For all allowed secretion reactions *j_EX_* ∈ {*Allowed secretions*}, their upper bounds 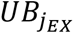 of exchange flux 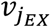 are set to an arbitrary large number *M* (i.e., 1e6 in this work). Nutrient uptake values and allowed secretory metabolites are recorded in the https://github.com/hvdinh16/iHME/resources.

### Metabolic task formulation

Metabolic task^44,48^ was utilized over biomass objective function (BOF) (i.e., lumped objective of elementary metabolic tasks)^92^ as the optimization objective for metabolic modeling. There are several compounding technical reasons for this choice. First, qualitative evaluation can be performed with metabolic task objective (i.e., successful task execution or not), unlike BOF whose exact biomass coefficients need to be specified. Second, biochemical measurements are currently incomplete across metabolites in BOF and across different growth condition contexts. Third, effects of gene and pathway perturbation are more traceable with metabolic task than with BOF (i.e., as a lumped objective).

To establish a metabolic task panel we first consulted the Agren et al., 2014’s core task^44^ panel (i.e., referred to as P1) which has been commonly used for essential gene evaluation. Instead of directly using the P1 core task panel, we created our own panel (i.e., referred to as P2) that has simpler modeling implementation. To compare, an original core P1 task requires extensive model reconfigurations for model-task interfacing, including adding unique sources, adding unique sinks, and adding unique conversions. For example, the core task “ATP de novo synthesis” has the following configurations: (i) sourcing of oxygen, glucose, ammonium, and phosphate; (ii) sinking of water, CO_2_, and ATP; and (iii) conversion of ATP + H2O <=> ADP + Pi. All of these were not part of the original model (i.e., not exchange reactions, which exist separately) and have to be added to the model for task evaluation and then removed. In our implementation P2, a simple demand (i.e., sink) reaction is added to the model (i.e., as ATP → PPi). We combine into a panel (P2) metabolic tasks created by three methods: (i) Adding demand reaction for metabolites in BOF, (ii) adding demand reaction of pathway endpoint containing essential genes, and (iii) adding source or conversion reaction for clearance and maintenance objectives (see Supplementary Fig. S1 for illustrative examples). The metabolic task panel is provided in Supplementary Table 1. Details on task matching is provided in https://github.com/hvdinh16/CMDep/blob/main/input/metTask_match_toAgren2014.xlsx.

### Context-specific metabolic network reconstruction

The initial INIT algorithm^30^ was formulated by Agren et al., 2012. Here, we adopted and modified the INIT algorithm for various input data (i.e., network, metabolic task, gene expression, and nutrient availability) compatibility. INIT works by maximizing the number of reactions associated with highly expressed genes while doing the opposite for lowly expressed genes.

First, the expression score *E_j_* for reaction *j* were calculated as a function of gene expression levels *E_g_* of all associated genes *g* ∈ *G_j_*, as follows:

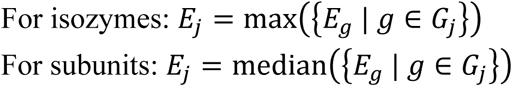

For complex gene-reaction rules, the expression score was compiled inside-out following the boolean logic. In short, boolean logic tree was reconstructed (i.e., executed via COBRApy and SymPy packages) and expression score of the branches of the tree are compiled from inside-out until a single value is obtained for the whole logic tree.

The formula for reaction score links potential of metabolic activities to major isozyme and to median expression of protein subunits. While enzyme complex formation is limited by the least abundant subunit, median is a better choice in general since subunit observation could be incomplete due to a lack of sequencing depth (i.e., making the least abundant subunit level zero). For enzyme complex with subunit isoforms of different genes, *w_j_* is sequentially calculated following the order of operation from the gene-reaction rule in the genome-scale metabolic reconstruction. For spontaneous, exchange, and intracellular demand (e.g., for a vitamin) reactions, *w_j_* = 0.01. For other non-gene associated reactions (i.e., belong to a hypothetical pathway), *w_j_* = −1. Then, the reaction score *w_j_* for reaction *j* (relative to the *E_cutoff_* is cutoff level of gene expression) was calculated as follows:

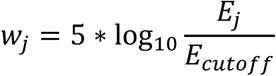

The version of INIT algorithm used in this work is as follows:

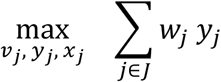

subject to

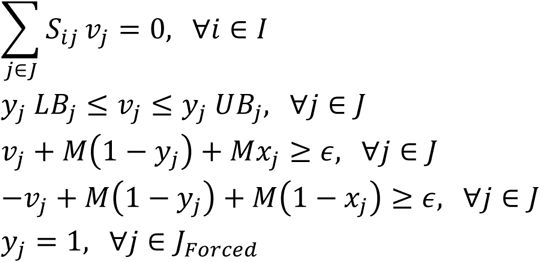

where *S_ij_* is stochiometric coefficient of metabolite *i* in reaction *j* and *I* and *J* are the sets of metabolites and reactions, respectively. *v_j_*, *y_j_*, and *x_j_* are flux variable, binary variable for reaction inclusion, and binary variable for forcing minimal flux in positive or negative flux direction, respectively. *LB_j_* and *UB_j_* are reaction lower and upper bounds, respectively. *LB_j_* is set to zero for an irreversible reaction and a large negative number −*M* otherwise. *UB_j_* is set to a large positive number *M*. *M* was set to 5000. *y_j_* = 1 will allow reaction *j* to carry flux and forcing the flux variable *v_j_* to be at minimal level of *ε* (set at *ε* = 1). *x_j_* is needed to handle the flux value in positive and negative direction.

Several reactions in the set *J_Forced_* are forced to be on by default. These include the demand, source, and conversion reactions representing metabolic tasks (i.e., task protection). Several other pathways that are inconsistently retrieved despite having essential genes have to be forced, including nitrogen balance (glutamine, glutamate, and ammonium uptakes and secretions), essential nutrient exchange reactions, glycolysis, pentose phosphate pathway, NADH shuttle (aspartate – glutamate and glyceraldehyde 3-phosphate – DHAP), TCA cycle, oxidative phosphorylation, *de novo* purine and pyrimidine synthesis, nucleotide synthesis, and cytosolic one-carbon metabolism.

If flux constraints are imposed (i.e., by setting in *LB_j_* and *UB_j_*), gapfill algorithm needs to be performed. This is because INIT only ensures flux balancing at minimal level but not physiological level at which metabolic overflow could occur. For example, glycolytic flux can theoretically be balanced with only respiration or fermentation, but imposing physiological glucose uptake and byproduct secretion constraints requires a network with all three pathways. The gapfill algorithm is performed as follows:

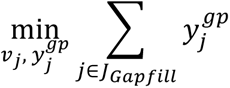

subject to

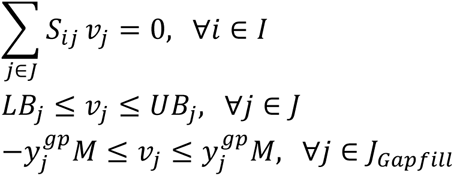

where *J_Gapfill_* is the set of reactions that are not included after performing INIT and become gapfilling candidates, and 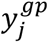 is binary variable to specifically using reaction *j* for gap-filling or not.

### Context-specific gene-reaction rule reconstruction

After network reconstruction, gene-reaction rules were also rebuilt (for all reactions being selected, or *y_j_* = 1) to eliminate redundant isozymes. Associated genes with expression level *E_g_* lower than reaction score *w_j_* or cutoff threshold *E_cutoff_* were removed from the gene-reaction rules, except for cases where it is absolutely necessary to retain the gene without breaking the gene-reaction rules (to be hereby explained).

The algorithm to rebuild gene-reaction rules is as follows. First, the boolean logic tree is reconstructed from the gene-reaction rules (i.e., executed via COBRApy and SymPy packages). Boolean logic components (of genes) in parentheses are branches of trees and are solved first, meaning that the logic is solved inside-out. Per branch, the expression score of a branch is compiled from *E_g_* of genes, where median value was assigned from subunits (i.e., logic “and”) and maximal value was assigned from paralogs (i.e., logic “or”). The rules for trimming of genes are as follow:

i. If the branch score is higher than *E_cutoff_* and genes are in the logic “or”, genes with *E_g_* < *E_cutoff_* are trimmed.
ii. If the branch score lower than *E_cutoff_* and genes are in the logic “or”, meaning that even the most expressed paralog (or isozyme) is below the cutoff, all genes are trimmed except for the gene with highest *E_g_*.
iii. Genes that are in the logic “and” are retained regardless of their *E_g_* values.

### Gene knockout simulation

Gene knockout simulation is performed for the context-specific network with reaction subset *J^c^* ⊂

*J* and metabolite subset *I^c^* ⊂ *I* and the context-specific gene-reaction rules. If the knockout of gene

*g* is sufficient to disable reaction(s) *j* ∈ *J^c^*^,*KO*^per gene-reaction rule, the reaction flux(es) is set to zero in the following FBA simulation that is performed per metabolic task (i.e., for all (*j* = *task*) ∈ *T* ⊂ *J^c^*, qualitatively tracked by flux variable *v*_(*j*=*task*)_):

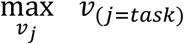

subject to

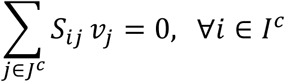

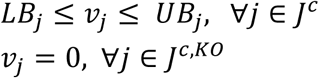

If ∃*task* ∈ *T*, *v_task_* = 0, or in words, there are at least a task that is disrupted by gene knockout, the gene *g* is considered essential.

### Model configuration and simulation protocol

To model the context-specific metabolism, base model reconstruction is performed first, then context-specific reconstruction is performed second, and lastly flux balance analysis (i.e., including essential gene analysis). The base model adjustments and curations were performed in iterative cycles subject to the false positives and negatives identified in context-specific model simulation. Through successive iterative cycles, these false predictions were resolved using rationales mentioned in Section 4.1. Once the final version of the base model was generated, transcriptional RNA-seq data (i.e., from DepMap^46^ and HPA^13^ database for cell line and tumor case studies, respectively) was prepared as a (genes x contexts) data table recording gene expression in transcript-per-million (TPM). Before simulation, user can configure inputs, include nutrient constraints, metabolic task panel, and task and pathway protection on/off. Setting options for cancer metabolism is available from this study, includes *in vitro* and *in vivo* nutrient constraints, proliferative task panel (i.e., 88 tasks), and essential task and pathway protection (i.e., see Section 4.4). Using the in-house developed reconstruction-and-analysis pipeline (i.e., implemented in Python programming and GAMS modeling languages), INIT network reconstruction, gene-reaction rule reconstruction, and essential gene analysis were performed with transcriptional data input. This resulted in context-specific models (i.e., one per context) and essential gene prediction (i.e., gene being essential per task per context). The software implementation of the pipeline is available at https://github.com/hvdinh16/INIT_iHME.

### Essential gene validation in cancer cell lines

From DepMap database (version 24Q4), RNA-seq and gene score data were collected for 1,103 cell lines for which both modalities were available. Using the pipeline-generated context-specific models and essential gene predictions were generated for cancer cell lines. Since the outcomes of prediction and observation are either essential or not, the prediction is classified into four types: true positive (TP), true negative (TN), false positive (FP), and false negative (FN). Outcome being positive indicates gene is essential. *TP*, *TN*, *FP*, and *FN* will be used in formula as counts of the respective prediction outcome. Recall is the ratio of *TP*/(*TP* + *FN*). Matthew’s Correlation Coefficient (MCC) is a balanced score calculated from the counts of all four prediction types, using the following formula:

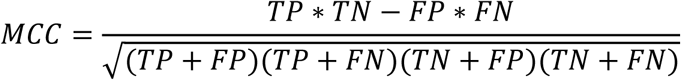

To identify successful predictions of context-specific essentiality, we create a control of essential gene prediction without RNA-seq data and thus the same base network was used for prediction for all cell lines. Here, the base network is still informed by nutrient constraints and metabolic tasks. Increase in MCC score, which arises from higher rates of TP and TN, is used to rank improvement in context-specific gene predictions with additional inputs of RNA-seq data.

For two groups, FDFT1-essential and non-essential cell lines, differential gene expression analysis was performed with two-tails t-test for metabolic gene subset (n = 1,430) with cutoffs of fold-change > 1.5 and FDR < 0.05. Differential DepMap gene score analysis was performed with one-tail t-test for metabolic gene subset (n = 1,430) with cutoff of FDR < 0.1. DepMap’s record of Oncotree classification for cell lines was used to further partition the essential and non-essential groups to the classification-specific components, and fold- and score changes were reported.

To examine whether DepMap gene score is significant for SREBP2-signaling cascade genes, differential score analysis was performed with one-tail t-test for all genes (n = 17,916 genes) with cutoff of FDR = 0.1. SREBP2-signaling genes^93^ include INSIG1, INSIG2, SCAP, SREBF2, SAR1A, SAR1B, SEC23A, SEC23B, SEC24A, SEC24B, SEC24D, SEC13, SEC31A, SEC31B, MBTPS1, MBTPS2, and PARQ3.

### Selection of priority genes for tumors

From Human Protein Atlas (HPA) database, the tumor transcriptional RNA-seq data (i.e., which is parts of Human Pathology Atlas project^13^) were retrieved (https://www.proteinatlas.org/about/download). HPA contains 8,384 samples, 6,918 of which were from TCGA. Metabolic network (reaction presence), gene presence, and essential gene information were generated. Essential frequency was calculated across 8,384 samples.

For tumor vs. normal tissue differential expression, raw count data were retrieved from TCGA database via UCSC Xena Browser (https://xenabrowser.net/datapages/), for paired tumor and tumor-adjacent tissue samples (from the same patient). Data was available for 17 out of 21 TCGA subtypes, which excluded GBM, SKCM, OV, and TGCT. Per TCGA subtype, raw counts of normal and tumor samples were first normalized by trimmed mean of M-values (TMM) method, and then paired t-test was performed. For GBM, TPM data for paired normal tissue and tumor was retrieved from GEO database deposited at GSE165595, described in Hwang et al., 2022^ref. 94^. Paired t-test was performed on uncorrected TPM data. For OV, pre-normalized data (by DESeq2) for unpaired normal and tumor samples was retrieved from supplementary materials in Carvalho et al. 2022^ref. 78^. Then, DESeq2 was performed on pre-normalized data to generate statistical p-values. For SKCM, raw microarray data for unpaired normal and tumor samples from Talantov et al., 2005^ref. 95^ was processed by the built-in GEO2R service of GEO database. Expression (in log2(count)) and p-value from statistical test performed by GEO2R were retrieved. All p-values were adjusted by Benjamini-Hochberg procedure to generate false discovery rate (FDR). Priority genes were selected by log2-fold change > 1 and FDR < 0.05.

For comparison of expression level between different genes in the same sample, TPM expression data from TCGA samples recorded in HPA dataset were used.

Differential DepMap gene score analysis was performed exactly for the designated gene (i.e., not multiple tests) using one-tail t-test with the cutoff of p-value < 0.05. The comparison was done for predicted essential vs. predicted non-essential cell line groups.

### *In silico* lethal drug prediction

Experimental PRISM secondary screening^6^ results were downloaded from DepMap portal^14^, dataset “PRISM Repurposing Secondary Screen”, version 19Q4. The overlapping 402 cell lines between DepMap RNA-seq and PRISM datasets, and 145 drugs targeting metabolic pathways in *i*HME model were evaluated in model simulation. Lethal drug incidence (per cell line per drug) was declared if the relative area under curve (AUC) of the response curve is below 0.8. Model prediction returned essential if at least a metabolic task is disrupted by turning off reactions inhibited by drugs. Inference from DepMap^46^ returned essential if drug targeting at least an enzyme that correspond to the gene with gene score below −0.5. For either model prediction or DepMap inference, the prediction outcomes are either true positive, true negative, false positive, or false negative; where positive indicates growth reduction of cells (i.e., lethal drug).

### Gene essentiality prediction with machine learning models

To benchmark our model against current machine learning models, we predicted gene scores using Exp-DeepDEP and DeepMeta using the same DepMap 24Q4 expression dataset as inputs. For Exp-DeepDEP, input data were prepared by the Prep4DeepDEP package (https://github.com/ChenLabGCCRI/Prep4DeepDEP, ‘Prediction’ mode, creating an input layer of 6,016 genes. Gene scores for 1,298 genes (in the output layer) were then predicted by the pre-trained model (https://codeocean.com/capsule/7914207/tree/v1). Gene was classified as predicted essential if predicted gene score was below −0.5. For DeepMeta (i.e., https://github.com/XSLiuLab/DeepMeta/tree/main), the input data were prepared from expression dataset with the R functions ‘PreEnzymeNet’ and ‘PreDiffExp’, and then essential reaction predictions were performed with the pretrained model. DeepMeta output was binary essential reaction predictions. Essential gene prediction was generated from reaction prediction by gene-reaction rule mapping. Genes that were in the base metabolic network but not context-specific networks used in DeepMeta were considered non-essential.

### Software and data availability

Software-based analyses were centralized on the Python (3.10.13) platform. Data processing was performed with pandas (2.3.3) and numpy (2.2.6). Mixed-integer linear optimization necessary for context-specific model reconstruction was solved by CPLEX solver (22.1.1) on General Algebraic Modeling System (GAMS) software (46.5.0). Linear programming necessary for flux balance analysis and gene essentiality prediction was solved by COBRApy (0.29.0) and CPLEX solver (22.1.1). Boolean gene-reaction rules were processed with sympy (0.13.2). Functions necessary for differential gene expression analysis were performed with rnanorm (2.1.0) and pydeseq2 (0.5.2). Statistical tests were performed with scipy (1.15.3). UMAP was performed by umap-learn (0.5.7). Visualization was performed by matplotlib (3.10.3) and seaborn (0.13.2). Program with graphic user interface was first developed in R (4.4.2) with rshiny (1.13.0) and further extended by AI model (Claude Code with Opus 4.6).

The base genome-scale model and associated files are deposited at https://github.com/hvdinh16/iHME_model. Cancer model libraries (1,013 cell lines and 8,384 tumor samples) are deposited at https://github.com/hvdinh16/iHME_model/tree/main/projects. The software implementation of the reconstruction algorithm is available at https://github.com/hvdinh16/INIT_iHME. Software scripts of reconstruction and analysis of results for cancer metabolic dependency are deposited at https://github.com/hvdinh16/CMDep. The CMDep repository plus heavy files are deposited at a Zenodo repository, https://doi.org/10.5281/zenodo.20804365. Program with graphic user interface for re-analysis of results are deposited at https://github.com/jiazhenz026/iHME_GUI.

## Supporting information

Supplementary Table - cell line results

Supplementary Table - tumor sample results

## Acknowledgements

The authors want to thank Caroline Bartman for discussions on metabolic modeling, and Sarah Cherkaoui for the suggestion on data analysis. This work was funded in part by the National Science Foundation’s Science and Technology Center for Engineering MechanoBiology, grant CMMI: 15-48571, to Y. Shen.

