## Supplementary Document for "A new genome-scale model enables prediction of cancer metabolic dependencies"

### List of materials

*List of Supplementary Figures*

Supplementary Figures and descriptions are included in this document.

Figure S1. Metabolic task formulation and task types
Figure S2. Method for prediction of human cancer essential gene with genome-scale model
Figure S3. Summary of *i*HME model reconstruction workflow
Figure S4. Reaction counts for *i*HME and Recon models
Figure S5. Prediction of lethal drug administration to cell lines
Figure S6. Gene essentiality prediction statistics by pathway
Figure S7. Prediction score comparison between models and gene subsets
Figure S8. Analysis of accurately predicted context-specific essential isozymes
Figure S9. Differential DepMap gene knockout score analysis for SREBP2 signaling genes
Figure S10. CDS2 dependency of skin tumor

*List of Supplementary Tables*

Supplementary Tables are provided separately in spreadsheets.

Table 1. Metabolic tasks
Table 2. Reactions of *i*HME model
Table 3. Metabolites of *i*HME model
Table 4. Genes of *i*HME model
Table 5. Prediction outcomes of context-specific and context-free models for human cancer cell lines
Table 6. Characteristics of priority genes in human tumor

*List of Metabolic Analysis Resources*

Resources and software implementations are provided in github repositories. Itemized resources, descriptions, and instructions are provided below.

iHME_model – Repository of *i*HME modeling resources
Link: https://github.com/hvdinh16/iHME_model
INIT_iHME – Repository of INIT software implementation
Link: https://github.com/hvdinh16/INIT_iHME
CMDep – Repository of cancer metabolic dependency analysis

Link: https://github.com/hvdinh16/CMDep

Zenodo link (including heavy files): https://doi.org/10.5281/zenodo.20804365

iHME_GUI – Repository of graphic user interface for cancer metabolic reconstruction

Link: https://github.com/jiazhenz026/iHME_GUI

### Supplementary Figures

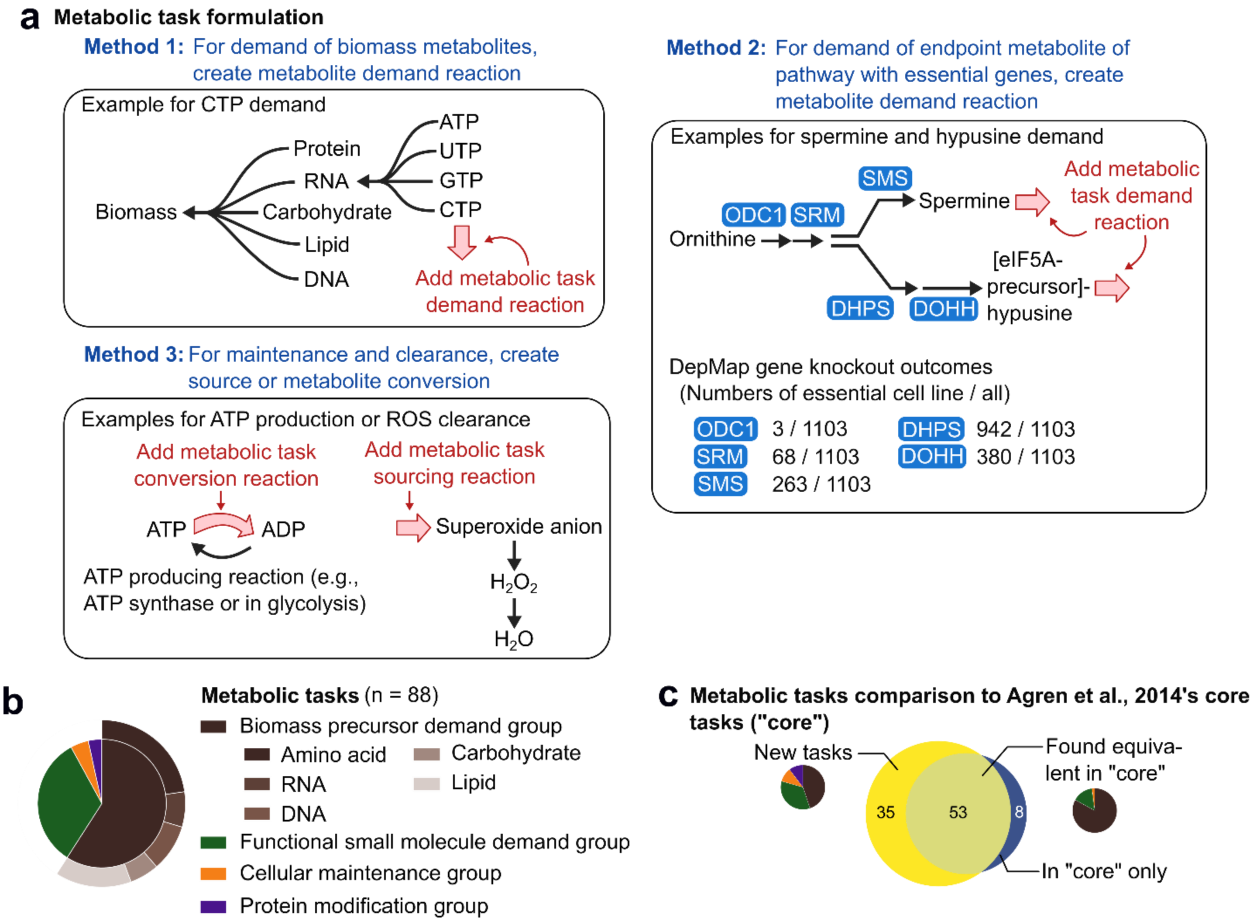


**Figure S1. Metabolic task formulation and task types**.

(**a**) Visualizations of three methods for metabolic task formulation.

(**b**) Summary of metabolic task panel (n = 88).

(**c**) Comparison of this panel to the core panel in Agren et al., 2014 (https://doi.org/10.1002/msb.145122). "Found equivalent in core" indicates that a task in this panel can be matched to another one from "core". "New tasks" indicates that a task in this panel cannot be matched. "In core only" indicates that a task in "core" panel cannot be matched.

These new tasks provide the basis for the model to capture more metabolic essential genes.

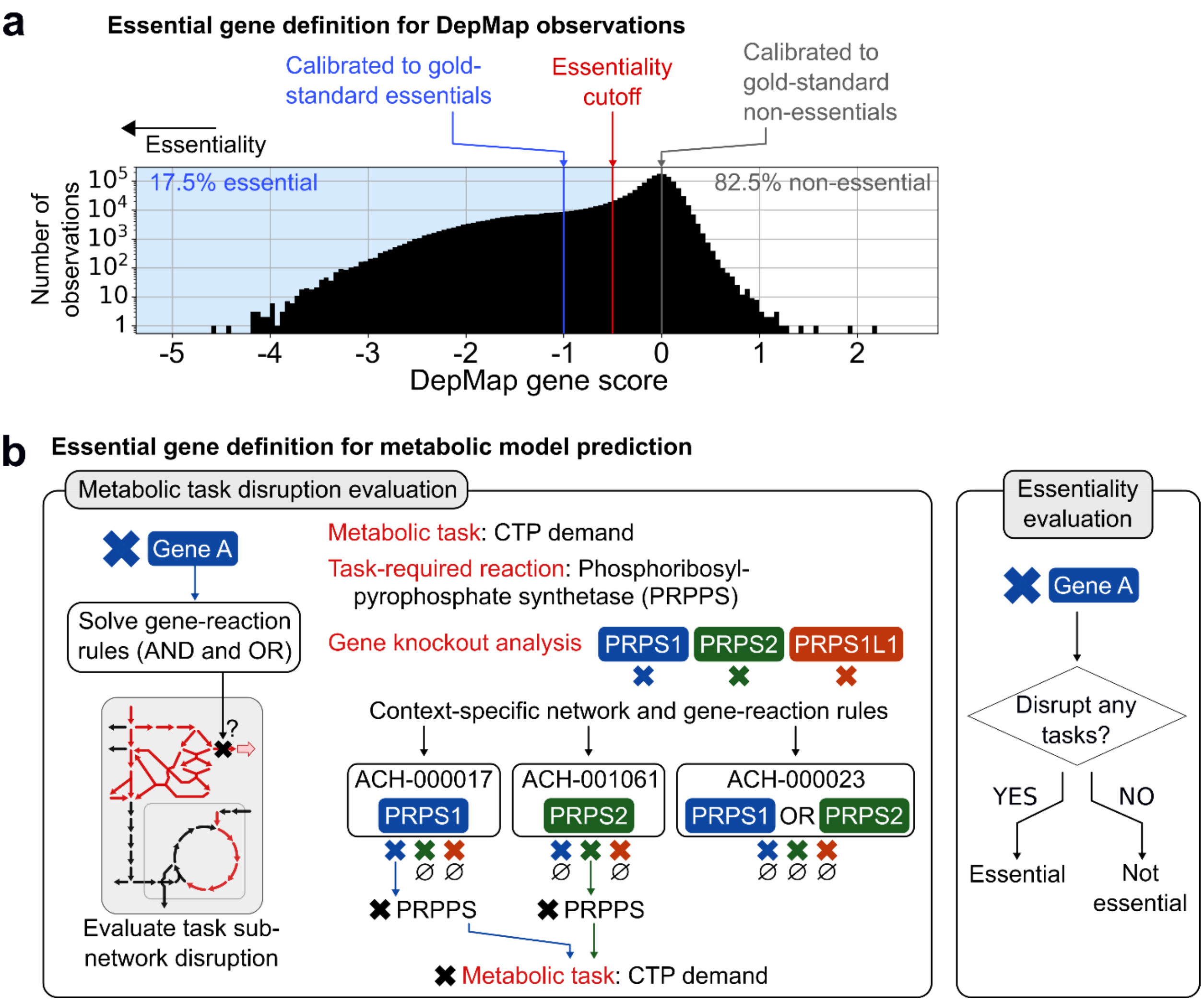


**Figure S2. Method for prediction of human cancer essential gene with genome-scale model.** (**a**) Histogram of DepMap gene scores cumulative across metabolic genes. In total, there are 1.58 million data points across 1,103 cell lines and 1,490 genes. Since DepMap gene score is calibrated so that the average score of gold-standard essential genes is negative one and the average of gold-standard non-essential genes is zero, the midpoint minus 0.5 is selected to be the cutoff for declaring essential and non-essential genes. y-axis of histogram is in log-scale.

(**b**) Visualization of method to declare gene essentiality in prediction. First, the model checks whether disrupting gene leads to reaction disruption (i.e., no isozyme rescue). Second, the model checks whether turning-off reaction leads to task disruption (i.e., reaction is essential and there is no alternative pathway), one-task-at-a-time. Third, the model declares that gene is essential if reaction turn-off disrupts at least a task in the panel. Altogether, experimental data and predictions will be used to classify whether predicted essentiality (per gene per cell line) is true positive, true negative, false positive, or false negative.

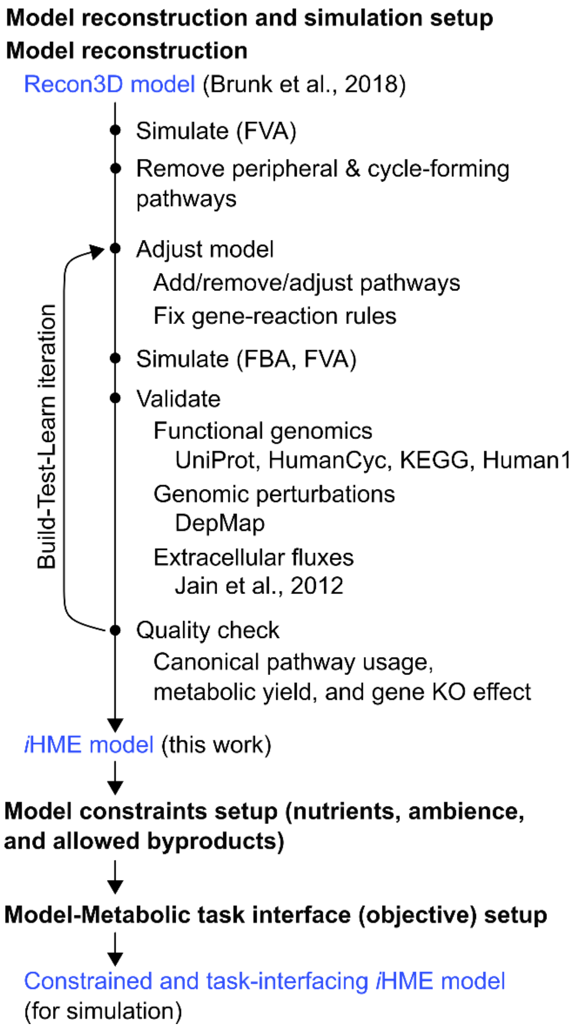


**Figure S3. Summary of *i*HME model reconstruction workflow**. This workflow matches the description provided in the method. To summarize, first, model was pruned by removing peripheral and cycle-forming pathways through analyzing by FVA. Second, in an iteration, model was adjusted, (re)simulated, and validated with biochemical database information and biochemical data. Quality check was performed to decide whether another iteration was needed or the fixing process could be terminated. Third, once the model passed quality check, it was integrated with nutrient constraints and metabolic tasks to become simulation-ready.

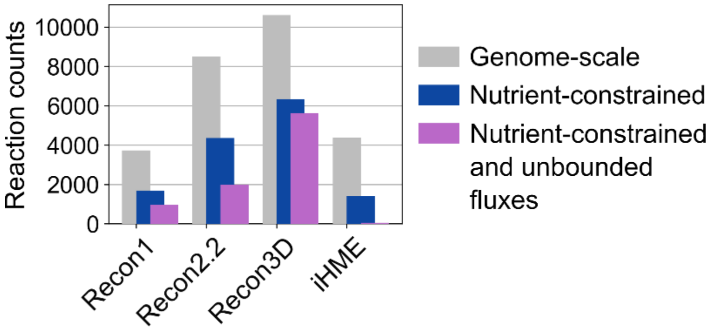


**Figure S4. Reaction counts for *i*HME and Recon models**. "Genome-scale refers" to the raw models without dropping out blocked reactions identified by FVA. "Nutrient-constrained" refers to the model with only reactions active under constraints retained. "Nutrient-constrained and unbounded fluxes" refers to the subset of "nutrient-constrained" reactions with flux values touching the large-number upper and lower bounds (i.e., nutrient allowed of one unit, bounds of 1000 units).

This figure demonstrates that Recon and *i*HME models did not converge to the same network size under the same nutrient constraints, supporting the claim that the size difference between Recon3D and *i*HME models cannot be explained by comprehensiveness for alternative nutrients. In addition, many thermodynamically infeasible cycles (i.e., unbounded fluxes) were found in Recon models.

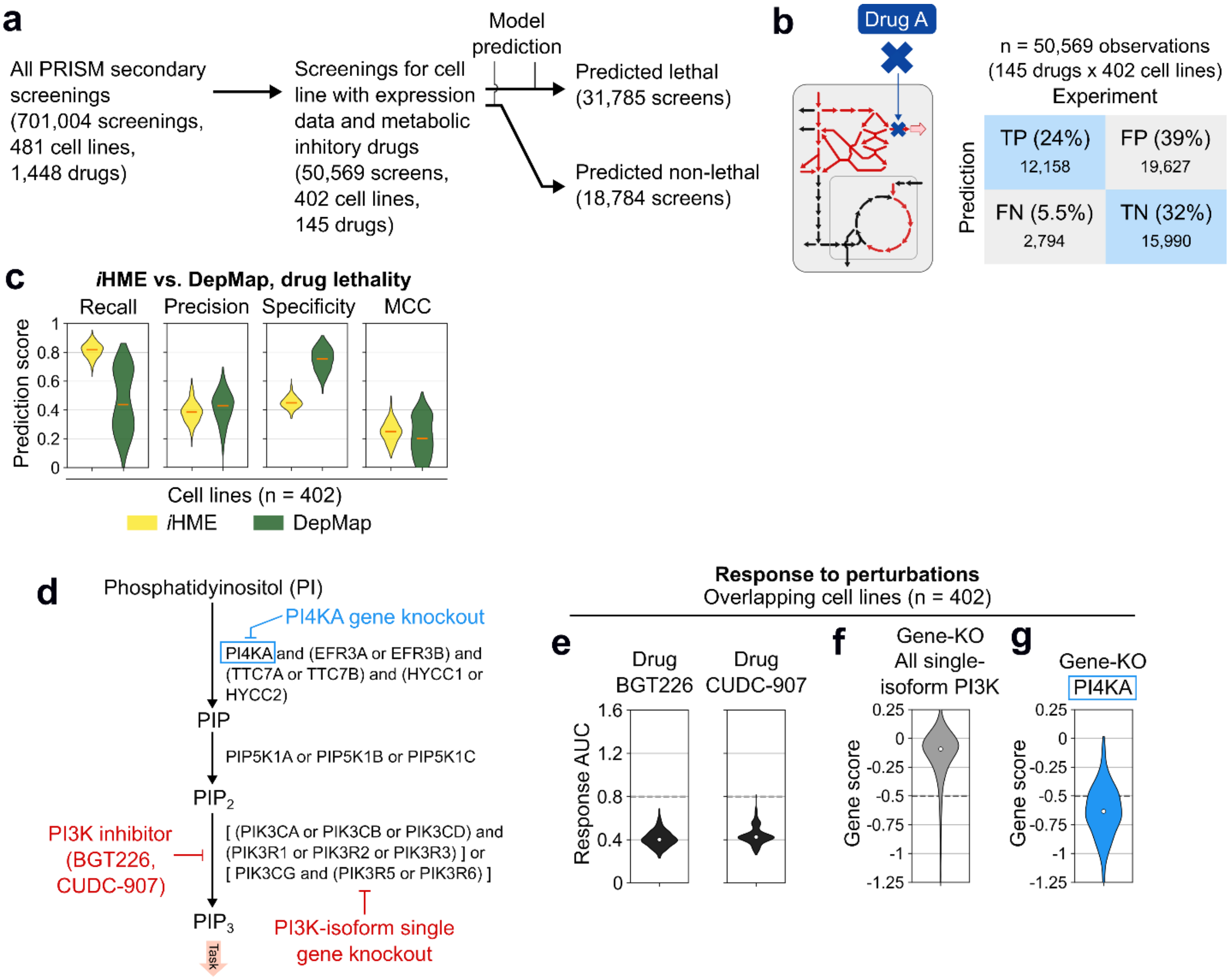


**Figure S5. Prediction of lethal drug administration to cell lines.**

(**a**) Selection of cell lines in reconstruction library and metabolic inhibitory drugs that can be simulated by the model.

(**b**) Statistics of two-by-two prediction-data matrix.

(**c**) Prediction scores distribution for cell lines (n = 402) using *i*HME model prediction or inference from DepMap. DepMap inference was performed by declaring drug is lethal if it disrupts the context-specific essential gene for a cell line. Orange line marks distribution medians.

(**d**) Schematics of gene knockout and PI3K inhibition in phosphatidylinositol (3,4,5)-triphosphate ($PIP_3$) synthesis pathway. This illustrates that inference from gene knockout will not account for cases where gene knockout is rescued by isozyme, but drug inhibition is not rescuable.

(**e**) Response AUC of two most effective PI3K inhibitory drugs, indicating PI3K inhibition is lethal.

(**f**) Gene knockout effects for all PI3K isoforms, showing no indication of PI3K being lethal.

(**g**) Gene knockout effect of PI4KA gene which has no isozyme, showing indicating that $PIP_3$ synthesis disruption is lethal.

This figure demonstrates the model applicability in predicting the effect of metabolic inhibition by drug.

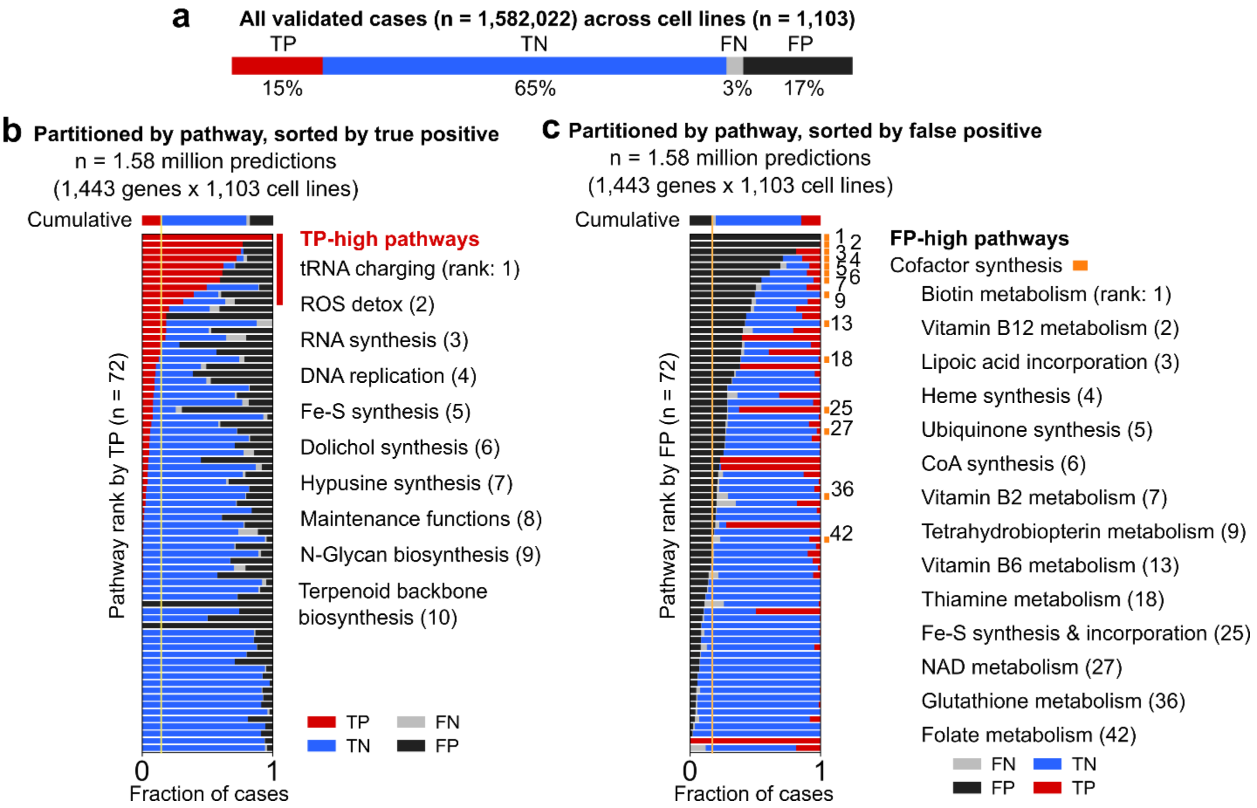


**Figure S6. Gene essentiality prediction statistics by pathway.**
(**a**) Cumulative statistics.
(**b**) Statistics by pathway, ranked by true positive fraction.
(**c**) Statistics by pathway, ranked by false positive fraction. In all plots, proportion of the four types in four colors indicates their relative fractions. In panel (**c**), cofactor synthesis pathways are marked with orange rectangles.
This figure illustrates that false positive has stronger association to cofactor synthesis rather than to other pathways.

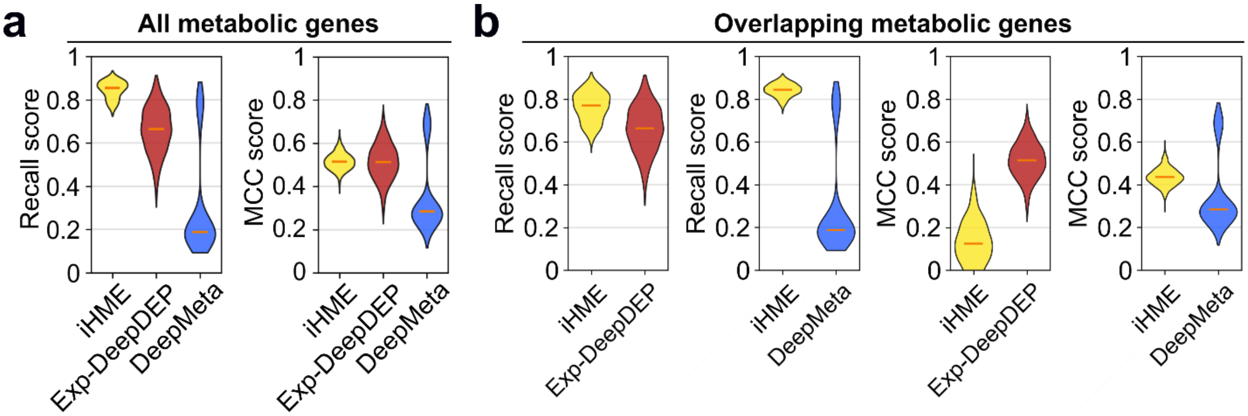


**Figure S7. Prediction score comparison between models and gene subsets**.
Recall and Matthew Correlation Coefficient (MCC) scores were compared for different models on different gene subsets. Here, *i*HME is the genome-scale mechanistic model of metabolism and Exp-DeepDEP and DeepMeta are deep learning models.
(**a**) For all metabolic genes (i.e., 1,487 genes in *i*HME, 180 genes in Exp-DeepDEP, and 485 genes in DeepMeta), for all cell lines, scores were calculated.
(**b**) For overlapping metabolic genes, *i*HME – Exp-DeepDEP (i.e., 180 overlapping genes) and *i*HME – DeepMeta (i.e., 485 overlapping genes) model pairs are evaluated separately. Scores were calculated for all cell lines.
Distributions are for scores across cell lines, n = 1,103 for iHME, 1,103 for Exp-DeepDEP, and 784 for DeepMeta due to its data selection. Orange line marks distribution median.

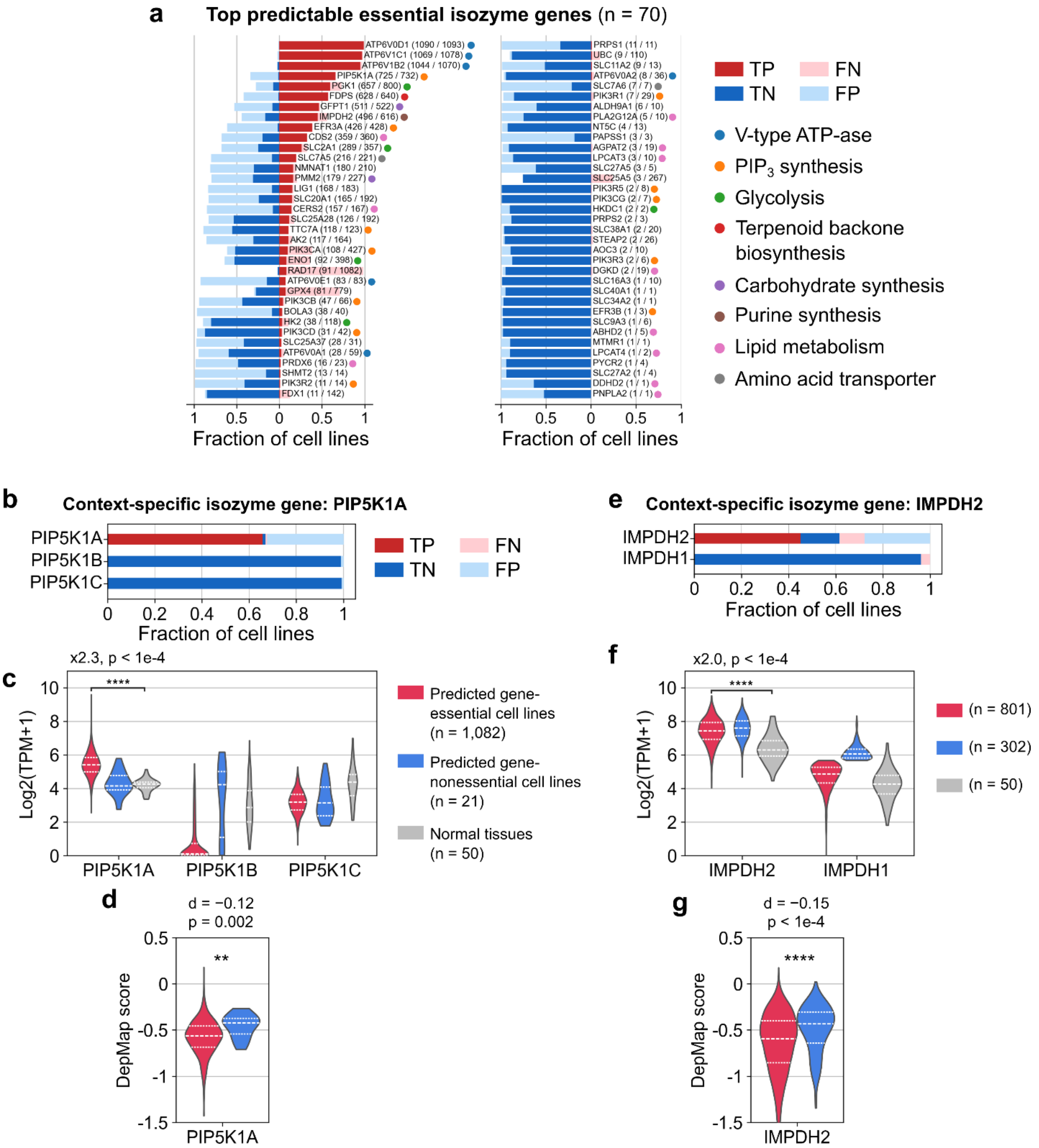


**Figure S8. Analysis of accurately predicted context-specific essential isozymes**.
(**a**) Detailed prediction statistics of different genes, with pathway marked by colored circles.
(**b,e**) Prediction statistics of isozymes for (**b**) PIP5K1A and (**e**) IMPDH2.
(**c,f**) Expression levels of different isozymes for (**c**) PIP5K1A and (**f**) IMPDH2.
(**d,g**) Difference in DepMap score between essential and non-essential groups for (**d**) PIP5K1A and (**g**) IMPDH2.
Cancer cell lines expression data is from DepMap database. Normal tissue expression data is from the consensus expression dataset of the Human Protein Atlas.

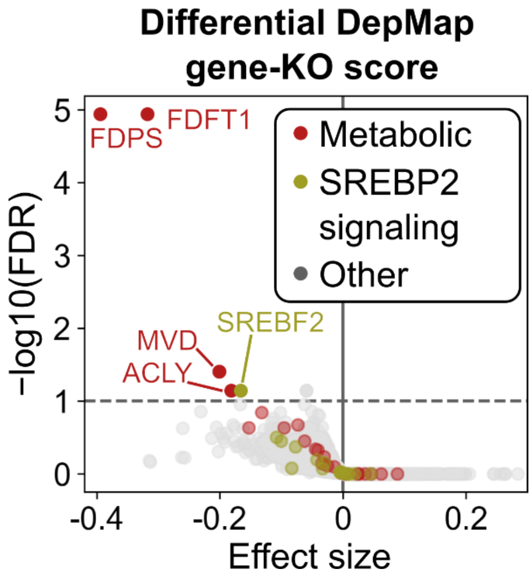


**Figure S9. Differential DepMap gene knockout score analysis for SREBP2 signaling genes**. Cholesterol synthesis is downstream of SREBP2 (i.e., a transcription factor, encoded by SREBF2 gene) signaling. The group of cell lines predicted by the model to depend on cholesterol synthesis (via FDFT1 essentiality) experience more severe growth reduction under knockout of cholesterol synthesis metabolic genes (denoted as red points). These cell lines also experience more severe growth reduction under SREBF2 knockout, suggesting an association between SREBP2 signaling dependency and cholesterol synthesis dependency in cancer cell lines.

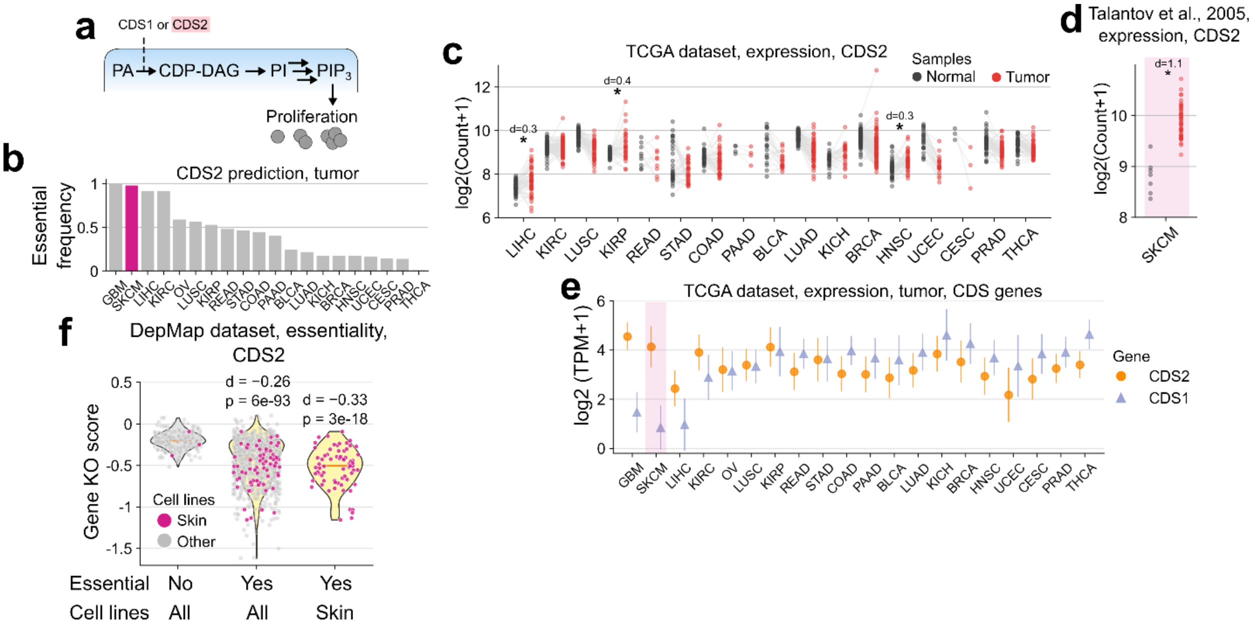


**Figure S10. CDS2 dependency of skin tumor.**

(**a**) CDP-diacylglycerol synthase (CDS) and $PIP_3$-synthesis pathway.

(**b**) Essential frequency in different TCGA tumor subtypes.

(**c**) Expression level in paired tumor and tumor-adjacent tissue samples from TCGA database. Expression is in log2(Count+1). Log2-fold change (d-value) and significance (*, if FDR < 0.05) from paired samples t-test are provided. N = 3 to 98 paired samples across TCGA tumor subtypes.

(**d**) Expression level in skin cell melanoma (SKCM) tumor and normal tissue samples from Talantov et al., 2005[ref.1]. Description is the same with (f) except for the usage of unpaired two-tail t-test. N = 45 and 7 for unpaired tumor and normal tissue samples, respectively.

(**e**) Expression level of glucose transporters in tumor. Expression is in log2(TPM+1). Mean and standard deviation values (across samples) are shown. N = 64 to 1,022 samples across TCGA tumor subtypes. HNSC and OV are highlighted.

(**f**) DepMap gene knockout scores across cell lines. Distributions of scores were plotted for groups of cell lines predicted by the model to be essential or non-essential and annotated to be from specific tissue of origins. One-tail t-tests were performed to compare scores of essential to those of non-essential groups.

**Metabolic Analysis Resources**

*iHME_model repository*

Link: https://github.com/hvdinh16/iHME_model
Repository of base genome-scale model

- Base genome-scale model
  - Full version (with gaps)
  - All nutrient exchange reactions on (gaps removed)
  - *In vitro* serum nutrient constrained version (gaps removed)
  - *In vivo* human plasma constrained version (gaps removed)
  - COBRApy json (for Python) and COBRA matlab (for MATLAB) formats
- Context-specific model from projects
  - DepMap 1,103 cell lines
  - HPA 8,384 tumor samples (including 6,918 TCGA samples)
  - File system descriptions
    - Reaction presence matrix
    - Gene presence matrix
    - Gene essentiality matrix
    - Lookup table – Sample ID and simple annotations
    - Parts – Detailed mechanism of gene essentiality mapped to exact metabolic tasks
- Resources
  - Biomass reaction information
  - Flux capacity information
  - Metabolic network visualizations by Escher (https://escher.github.io/)
- Scripts
  - Extract context-specific model from base model
  - Load flux constraints to model

*INIT_iHME repository*

Link: https://github.com/hvdinh16/INIT_iHME
Repository of software implementation of INIT algorithm for context-specific reconstruction

- Input
  - Flux capacity constraints information
  - Metabolic task objective testing settings
  - INIT optimization-specific settings
    - demand_reactions_allow
      Specific material sinks in the form of demand reactions. If new metabolic task is added, this file needs to be modified)

- essential_nutrients_min_amount
  Essential nutrients that need to be set to minimal amount. Essential nutrient set is a known fixed property of the model, and this will bypass missing or erroneous settings of zero essential nutrient uptake. These essential nutrients cannot be unconstrained because there are catabolic pathways for these nutrients.

- essential_nutrients_open
  Essential nutrients that need to be set to minimal amount. Essential nutrient set is a known fixed property of the model, and this will bypass missing or erroneous settings of zero essential nutrient uptake. These essential nutrients can be unconstrained because there are no catabolic pathways for these nutrients.

- exchange_allowed_generic/generic_vitroSerum/generic_humanPlasmaUpdate
  Allow exchange reactions. Membership of this set is limited to prevent unchecked secretion and uptake of random metabolites. This allows INIT algorithm to run even with incomplete description of extracellular flux profiles.

- exchange_media_ambiance
  Allow exchange reactions granted by being in ambient media, including in this work, but not limited to for other scenarios, oxygen, phosphate, iron, sulphate, proton, water, and CO2.

- exchange_secretion_allowed_default
  Allow default allowed secretions, including in this work, but not limited to for other scenarios, urea, CO, NO, uric acid, ammonium, and $CO_2$.

- simulation_settings_general
  Specialized optimization operation that does not fit into general above cases. This can be used in a versatile manner to make prediction that fit to a particular conditions in a case study.

- init
  List of forced sets of reactions to be on, including: NADH shuttle, OXPHOS, pentose phosphate pathway, TCA cycle, biomass components, *de novo* nucleotide synthesis, glycolysis, maintenance functions, one carbon

metabolism, protein modification, trace metabolites demand, nutrient uptake reactions, and miscellaneous special reactions. This folder just lists out different reaction sets. Setting which sets to be forced on has to be done in python workflow file.

- Model
  This is a location to place genome-scale model. Default files are the same to model files deposited at iHME_model
- Results
  This is a location to generate a project. This contains template to start a context-specific reconstruction project.
- Scripts
  Directory contains Python and GAMS scripts for context-specific model reconstruction.

*CMDep repository*

GitHub link: https://github.com/hvdinh16/CMDep
Zenodo link (including heavy files): https://doi.org/10.5281/zenodo.20804365
Repository of the modeling and computational analyses for cancer metabolic dependency project

- Input
  - Modeling, software, and data input
  - INIT software (as in INIT_iHME repository)
  - Formatted expression data
  - Formatted PRISM secondary drug screening data
- Model
  - Base genome-scale model (as in iHME_model repository)
- Scripts
  - Python and GAMS scripts (as in INIT_iHME repository)
- Results (context-specific reconstruction results)
  - DepMap_CCLEset: 1,103 cell lines. Drug lethality simulation results were also in here.
  - HPA_tumor: 8,384 tumor samples
  - reconTimeBenchmark: iHME vs. Recon models
  - DepMap_fullGenomeScale: genome-scale, without nutrient constraints, not context-specific
  - DepMap_nutsCon: with nutrient constraints, not context-specific
  - DepMap_panCancer: with nutrient constraints, only with reaction recorded in at least a context-specific cancer cell line model, not context-specific
- Analysis (in-depth analyses and follow-ups)
  - allCancerContext: analysis of tumor samples

- compare_Human1_Human2: comparing *i*HME results to Human1 and Human2 results
- compare_MLModels: comparing *i*HME results to deep learning model results
- compare_Recons: comparing *i*HME to Recon models
- context_specific: analysis of cancer cell lines
- DepMap_score: Visualizing DepMap score distribution and cutoff
- drug_screening: analysis of lethal drugs
- model_identity: overlap anlaysis between *i*HME and Recon3D
- scoreOverConstraints: Recall score between context-specific and context-free models.
- stats_by_pathway: true positive and false positive statistics by pathways

*iHME_GUI visualization software and repository*

Link: https://github.com/jiazhenz026/iHME_GUI
Software is provided to assist with browsing the same metabolic dependency results from this work with graphic user interface. Software can be installed from installation package releases on GitHub repository (for Windows and MacOS-arm64).
